# The “LINC” in between Δ40p53-miRNA axis in the regulation of cellular homeostasis

**DOI:** 10.1101/2022.04.28.489891

**Authors:** Apala Pal, Pritam Kumar Ghosh, Saumitra Das

## Abstract

Previous research has shown that Δ40p53, the translational isoform of p53, can inhibit cell growth independently of p53 by regulating microRNAs. Here, we explored the role of Δ40p53 in regulating the long non-coding RNA-microRNA-cellular process axis, specifically focusing on LINC00176. Interestingly, LINC00176 levels were predominantly affected by the overexpression/stress-mediated induction and knockdown of Δ40p53 rather than p53 levels. Additional assays revealed that Δ40p53 transactivates LINC00176 transcriptionally and could also regulate its stability. RNA immunoprecipitation experiments revealed that LINC00176 sequesters several putative microRNA targets, which could further titrate several mRNA targets involved in different cellular processes. To understand the downstream effects of this regulation, we ectopically overexpressed and knocked down LINC00176 in HCT116 p53−/− (harboring only Δ40p53) cells, which affected their proliferation, cell viability, and expression of epithelial markers. Our results provide essential insights into the pivotal role of Δ40p53 in regulating the novel LINC00176 RNA-microRNA-mRNA axis independent of FL-p53 and in maintaining cellular homeostasis.

## INTRODUCTION

The tumor suppressor p53 is a master transcriptional regulator that regulates several critical events during cell fate determination, including transcriptional regulation of multiple genes involved in cell survival, cell cycle regulation, apoptosis, and metabolism (Somasundaram 2000). p53 has 12 isoforms (created by alternate promoters, splicing sites, and translation initiation sites) which have unique functions and can modulate p53 activity (Bourdon, Fernandes et al. 2005). The Δ40p53 isoform is the only translational isoform of p53. Our laboratory has shown that translation initiation from the first IRES (Internal Ribosomal Entry Site) of *TP53* generates full-length p53 protein, whereas translation initiation from the second IRES generates Δ40p53 (Ray, Grover et al. 2006). Various ITAFs (IRES *trans*-acting factors) are required for the IRES-mediated translation of the two isoforms under different stress conditions (Grover, Ray et al. 2008, Sharathchandra, Lal et al. 2012, Khan, Katoch et al. 2015). We have also shown how a protein and miRNA crosstalk at the TP53 3′-UTR influences the differential expression of p53 and Δ40p53 (Katoch, George et al. 2017).

These two isoforms have a robust affinity towards each other as both contain the oligomerization domain and can form homo- and hetero-tetramers (Ghosh, Stewart et al. 2004). Δ40p53 can both positively and negatively regulate p53 activity: it can alter the recruitment of p53 co-activators, increasing its activity (Candeias, Malbert-Colas et al. 2008), but it can negatively regulate the transactivation of p53 target genes, such as p21 (Powell, Hrstka et al. 2008). Δ40p53 also has p53-independent functions. It controls the switch from pluripotency to differentiation in somatic cells and helps maintain pluripotency in embryonic stem cells. It suppresses the Nanog and IGF receptors and maintains pluripotency in embryonic stem cells (Ungewitter and Scrable 2010). It also regulates proliferation and glucose homeostasis in mice (Hinault, Kawamori et al. 2011).

Moreover, it uniquely regulates apoptosis-related genes such as p53BP2 and TIAL1(Ohki, Kawase et al. 2007) and can also independently induce senescence (Maier, Gluba et al. 2004). In certain cancers, full-length p53 is insufficient to protect against tumorigenesis, and Δ40p53 is essential for mediating apoptosis through the selective activation of specific target genes (Phang, Othman et al. 2015). Recent observations also suggest a role for Δ40p53 as a tumor suppressor in hepatocellular carcinoma (Ota, Nakao et al. 2017). Our latest studies showed that Δ40p53 could regulate miR-186-5p independent of full-length p53 (Katoch, Tripathi et al. 2021). However, some studies have examined the functional role of Δ40p53; there has been little detailed investigation regarding intermediate molecules (such as RNAs or proteins) through which Δ40p53 exerts its influence on cells.

Some RNAs in the p53 response pathway are involved in tumor suppression (Hunten, Kaller et al. 2015). Among the cellular pool of RNAs, long non-coding RNAs (lncRNAs) have gained attention in cancer research because of their transactivation functions. Several lncRNAs crosstalk with p53, thereby regulating cellular processes, including cell proliferation, differentiation and development, chromosomal imprinting, and genomic stability (Paralkar and Weiss 2013, Knoll, Lodish et al. 2015, Chaudhary and Lal 2017). However, there have been no studies on the role of Δ40p53 in lncRNA regulation. LncRNAs are a heterogeneous class of non-coding RNAs >200 nucleotides in length with very low or no coding potential. They also function as competitive endogenous RNAs (ceRNAs) that sequester miRNAs and alter the expression of their downstream target genes (mRNAs) by competing for their common binding regions. The ceRNA network links the function of protein-encoding mRNAs to that of non-coding RNAs. Growing evidence indicates that such networks formed by lncRNA-RNA-mRNA interactions have implications for a broad spectrum of biological processes, including disease conditions such as carcinogenesis (Wang, Cho et al. 2019, Seo, Kim et al. 2020, Gao, Zhao et al. 2021, Wang, Wang et al. 2021). In cancer cells, several aberrantly expressed long non-coding RNAs (lncRNAs) with miRNA responsive elements (MREs) competitively sequester miRNAs and impair the interaction between miRNA and mRNA, thereby attenuating the repression of downstream mRNAs (Wang, Cho et al. 2019). Thus, this emerging network is crucial for diverse cellular processes such as cancer (Poliseno and Pandolfi 2015, Su, Wang et al. 2021).

Many lncRNAs comprising such networks that regulate cancer progression remain underexplored (Hu, Sood et al. 2018, Zhao, Zhang et al. 2021). LINC00176 is a novel Myc-targeted lncRNA that has not been extensively studied. It has been identified as a *THOC5* target gene that regulates apoptosis and cell growth. Moreover, the ability of LINC00176 to titrate microRNAs in hepatocellular carcinoma (HCC) indicates its potential involvement in differential ceRNA networks, thereby governing critical cellular processes (Tran, Kessler et al. 2018). The same study also suggested that the promoter for LINC00176 is putative and that its localization may vary in a cell context-specific manner (Tran, Kessler et al. 2018). Another study screened for the functions of lncRNAs using genome-scale CRISPR screening. They showed that, upon deletion of some regions of LINC00176, it acts as a tumor suppressor lncRNA (Zhu, Li et al. 2016). Recent studies have also shown that LINC00176 is involved in ovarian cancer progression by downregulating ceruloplasmin expression (Dai, Niu et al. 2020). These reports highlight the function of LINC00176, which could be differentially expressed in different cells based on promoter localization and binding of different transcription factors. Although 80 transcription factors have been reported to bind LINC00176, little is known about how this gene is regulated (Tran, Kessler et al. 2018).

The two major transcription factors that govern antagonistic functions in cells are Myc and p53 (Sachdeva and Mo 2009). Myc regulates thousands of genes involved in cell survival and proliferation (Dang, O’Donnell et al. 2006), whereas p53 and its isoforms are stress-responsive molecules that activate the elimination of severely damaged cells (Zilfou and Lowe 2009). The frequently imbalanced expression of c-Myc over p53 signifies that the regulatory interactions between c-Myc and p53 are paralyzed in cancer cells (Evan and Vousden 2001). p53 and its isoforms play a central role in most known cancer pathways (Vieler and Sanyal 2018). Hence, we were curious to investigate the regulatory effect of p53 and its isoforms on LINC00176. To begin the study on Δ40p53- or p53-mediated regulation of lncRNAs, we primarily focused on LINC00176 and its possible impact on the ceRNA network and, in turn, its influence on cell growth and proliferation.

## MATERIALS AND METHODS

### Bioinformatic Analysis

The starBase v2.0 database (https://starbase.sysu.edu.cn/starbase2/browseNcRNA.php) was used to identify potential miRNA targets of LINC00176 (Li, Liu et al. 2014). PROMO-ALGGEN has been used to predict transcription factor binding sites in *LINC00176* gene (http://alggen.lsi.upc.es/cgi-bin/promo_v3/promo/promoinit.cgi?dirDB=TF_8.3). miRTarBase has been used to identify mRNA targets of miRNAs (https://mirtarbase.cuhk.edu.cn/~miRTarBase/miRTarBase_2022/php/index.php)(Huang, Lin et al. 2022). dbDEMC2.0 has been used to perform a meta profiling of selected miRNAs (https://www.biosino.org/dbDEMC/help#help00) (Yang, Wu et al. 2017).

### Cell lines and transfections

We used three cell lines in the current study: H1299 (lung adenocarcinoma cell line lacking Δ40p53 or p53 HCT116 p53+/+ cells), colon carcinoma cell line harboring wild-type p53 (hereafter called HCT116+/+), and HCT116 p53−/− cells (colon carcinoma cell line harboring only Δ40p53 (hereafter called HCT116−/−). HCT116−/− is a commercially available cell line with a deletion in the first ATG region of p53 mRNA; therefore, the only protein expressed from the mRNA is Δ40p53 (being the only translational isoform) from the second ATG. These cells were maintained in DMEM (Sigma) with 10% Foetal Bovine Serum (GIBCO, Invitrogen). 70%-80% confluent monolayer of cells was transfected with various plasmid constructs using Lipofectamine 2000 (Invitrogen) and Turbofectamine in Opti-MEM (Invitrogen). The medium was replaced with DMEM (antibiotic) and 10% FBS 4 h later. The cells were harvested and processed as required at the desired time point. For the stress-induced experiments, the cells were treated with doxorubicin (2 µM) for 16 h, thapsigargin (0.1 µM, 1 µM) for 16 h, and DMEM minus Glucose medium for 8 h. For stability experiments, cells were transfected with siRNA directed to Δ40p53; post 40 h of transfection, cells were treated with Actinomycin D (1 mg/ml), and cells were harvested at 0 h, 4 h, and 8 h post-treatment.

### Plasmids and constructs

pGFP-hp-p53- 5’UTR-cDNA (14A): It expresses both p53FL and Δ40p53 (a generous gift from Dr. Robin Fahraeus, INSERM, France). pGFP-hp-p53-5’UTR (A135T)-cDNA which expresses onlyΔ40p53 and pGFP-hp-p53- 5’UTR (A251G/T252 C/G253T)-cDNA which expresses only full-length p53(p53FL) (Katoch, Tripathi et al. 2021). Two constructs were used for Tagged RNA Affinity Purification: Plasmid pMS2 and pMS2-GST. (Both the constructs were a generous gift from Je-Hyun Yoon and Myriam Gorospe Laboratory of Genetics and Genomics, National Institutes of Health, Baltimore, MD 21224, USA) (Yoon, Srikantan et al. 2012, Yoon and Gorospe 2016). LINC00176 (3495–4302) construct obtained in the study (a generous gift from Dr. T Tamura Institut fuer Biochemie, Hannover, Germany) has been used to generate the overexpression construct. LINC00176 overexpression construct was generated by cloning the insert into pMS2 plasmid between HindIII and EcoRI sites. 5’CCGGTGTCGTCTTGCAAACATTAAACTCGAGTTTAATGTTTGCAAGACGAC ATTTTTG-3’ sequence of shRNA was used to target LINC00176.

### siRNA transfections

HCT116+/+ and HCT116−/− cells were transfected with 30 nM si p53 RNA (IDT). 5’- AACCUCUUGGUGAACCUUAGUACCU-3’ is the p53 siRNA sequence directed against the 3’UTR of p53; therefore, it targets both p53 as well as Δ40p53. A non-specific siRNA (si Nsp) was used in the experiments as a control (Dharmacon). si-LINC00176 (5’- CUCGUUCUGUAGACUUGUU-3’) was used for partial silencing of LINC00176 (a generous gift from Dr. T Tamura, Institut fuer Biochemie, Hannover, Germany) (Tran, Kessler et al. 2018).

### Western Blot analysis

Protein concentrations of the extracts were assayed by Bradford (Bio-Rad), and equal amounts of cell extracts were separated by SDS 12% PAGE and transferred to nitrocellulose membrane (Bio-Rad). Samples were then analyzed by Western blotting using a rabbit-raised anti-p53 polyclonal antibody (CM1, a kind gift from Dr. Robin Fahraeus, INSERM, France, and Prof. J.C. Bourdon, of University of Dundee, UK), followed by secondary antibody (horseradish peroxidase-conjugated anti-mouse or anti-rabbit IgG; Sigma). Mouse-monoclonal anti-β-actin antibody (Sigma) was used as a control for equal loading of total cell extracts. 1801 antibody (Santa Cruz Biotechnology, Cat. No. SC-98) was used for pull-down of p53 and Δ40p53 in the Immunoprecipitation experiments. E-cadherin antibody (Santa Cruz Biotechnology, Cat. No. SC-8426), Slug antibody (Santa Cruz Biotechnology, Cat. No. SC-166476), Bip antibody (Abclonal, Cat. No. A4908), and GST Antibody (B-14) (Santa Cruz Biotechnology, Cat. No. SC-138) was used to detect E-cadherin, Slug, Bip and GST proteins respectively. SIRT1 Antibody (Abclonal, Cat. No. A0230), Bcl-2 Antibody (Abclonal, Cat, No. 19693) and Cyclin E1 Antibody (Abclonal, Cat. No. 14225) was used to detect SIRT1, Bcl-2 and Cyclin E1 proteins respectively.

### RNA isolation and Real-time PCR

According to the manufacturer’s protocol, total RNA was isolated from cells with TRI Reagent TM (Sigma). The isolated RNA was treated with 10 units of DNase I (Promega), extracted with acidic phenol-chloroform, and precipitated with 3 M sodium acetate (pH 5.2) and 2.5 volumes of absolute ethanol. RNA amount was quantified in Nano-spectrophotometer, and cDNAs were synthesized using specific reverse primers and MMLV RT (Revertaid ™ Thermo Scientific) at 42°C for 1 h, according to standard protocol. While 2-5 µg of total RNA has been used for checking the expression of mRNAs and lncRNAs, 50 ng of total RNA has been used to check the expression of miRNAs.

SYBR green Assay System was used for mRNA, lncRNA, and miRNA detection and quantification. We have used actin as an endogenous control for mRNA, lncRNA, and 5SrRNA as an endogenous control for miRNAs. The thermocycling conditions for SYBR green Assay system include 1 cycle of 95°C for 10 min, 40 cycles of 95°C for 15 s, 60°C for 30 s, and 72°C for 30 s (Applied Biosystems). 2^−ΔΔCt^ method algorithm was used to analyze the relative expression changes, where actin/5S served as an endogenous control. The fold change was calculated using 2^−ΔΔCt^. ΔCt = Ct (target gene) −Ct (endogenous control) and ΔΔCt = ΔCt target sample) −ΔCt (control sample). Melting curve analysis of every (q)PCR was conducted after each cycle.

### Chromatin Immunoprecipitation (ChIP)

ChIP was performed with Human colon carcinoma cell lines (HCT116+/+ and HCT116−/−). Five 60 mm dishes were seeded with 0.399 × 10^6^ cells (each) and were allowed to grow for 48 h. The cells were harvested for processing using Sim-pleChIP® Plus Enzymatic Chromatin from Cell Signalling Technologies as per the manufacturer’s protocol. Briefly, cells were fixed using formaldehyde for 10 minutes at room temperature. The reaction was stopped using 1X Glycine. The cells were then collected in ice-cold 1X PBS (Phosphate Buffer Saline). The cells were lysed in the lysis buffers provided in the kit, digested with Micrococcal Nuclease, and processed (sonicated) to acquire nuclear chromatin lysate according to the protocol. The chromatin is then incubated with specific antibodies in the cyclomixer overnight at 4°C. Pull-down was performed using a p53 or Δ40p53 specific antibody. Magnetic protein-G beads supplied in the kit were then added to the antibody-bound lysate mixture and incubated in cyclomixer for 2 h at 4°C. The beads were then separated using a magnetic separator and washed with the wash buffer provided in the kit. Before the elution step, 20% of the beads were kept for western blot analysis to confirm equal pull-down of the protein. Elution was carried out in the elution buffer provided in the kit at 55°C for 30 minutes. It was followed by Proteinase K and 1% SDS treatment for 30minutes-2 h, followed by phenol-chloroform-based DNA extraction. The DNA was precipitated using 1/10th volume of 3M Sodium Acetate (pH 5.2) and 2.5 volume absolute ethanol at −20°C overnight.

### Copy Number Calculation

All miRNAs and LINC00176 RNAs were synthesized by *in vitro* transcription cDNA preparation and real-time PCR were performed using different increasing concentrations of RNA. The Ct values were obtained for each cDNA amount. The amounts were converted to Log2 of copy number, and a standard curve was plotted with Ct value on the Y-axis and Log2 (Copy Number) on the X-axis. Using the points obtained from the graph, a slope was generated, which was used to calculate the endogenous copy number of the miRNAs and LINC00176.

### Tagged RNA Affinity Pull down Assay (TRAP Assay)

The method is based on adding MS2 RNA hairpin loops to a target RNA of interest, followed by co-expression of the MS2-tagged RNA together with the protein MS2 (which recognizes the MS2 RNA elements) fused to an affinity tag (Yoon, Srikantan et al. 2012). After purification of the MS2 RNP complex, the miRNAs present in the complex are identified. In this study, we have tagged the LINC00176 with MS2 hairpins and have co-expressed it in Human colon carcinoma cells (HCT116−/−) along with the chimeric protein MS2-GST (glutathione S-transferase). After affinity purification using glutathione SH beads, the microRNAs present in the RNP complex

### Cell fractionation

Approximately 1×10^6^ cells (HCT116−/−) were washed with ice-cold PBS, resuspended in 175µl of Cytoplasmic lysis buffer (50mM Tris [pH 8.0], 140 mM NaCl, 1.5 mM MgCl2 and 0.5% NP-40) substituted with RNase Inhibitor (Thermo Fisher) and incubated on ice for 5mins. Cytoplasmic fraction was separated as supernatant by centrifugation at 500g at 4°C/4mins. The remaining nuclear pellet was resuspended in cold PBS and pelleted with high-speed centrifugation (5000rpm/1min). According to the manufacturer’s protocol, RNA was extracted from both fractions using TRI Reagent TM (Sigma). The relative transcript levels were determined in each fraction by normalization with their total levels.

### MTT Assay

HCT116−/− cells were transfected with either LINC00176 o/e or sh-LINC00176 construct 36 h post-transfection; these cells were re-seeded in 96 well plates. After 20 h of re-seeding, 10 µl of MTT (50 mg/ml) reagent was added, and cells were incubated for 3-4 h, following which the supernatant was removed, and cells were dissolved in 200µl of DMSO. Cell viability was measured as a spectrophotometry reading of the amount of MTT incorporated in the cell at 560 nm. The reading taken on the first day is considered Day 0. Similarly, the reading was measured on Day 2 and Day 4 post-re-seeding to check the effect on cell viability.

### BrdU cell proliferation assay

BrdU cell proliferation assay was performed as described in BrdU Cell Proliferation ELISA Kit (ab126556). Briefly, HCT116−/− cells were transfected with si-LINC00176. 36 h post-transfection, these cells were re-seeded in 96 well plates. After 12 h of re-seeding, BrdU solution was added, and cells were incubated for 12 h, followed by the protocol mentioned in the manual.

### Statistical analysis

All the experiments have been performed in three independent biological replicates (n=3). The data were expressed as mean ± SD. Statistical significance was determined using a two-sided Student’s t-test. The criterion for statistical significance was *p≤*0.05 (*) or *p≤*0.01 (**) or p*≤* 0.001(***)

## RESULTS

### Selection and validation of LINC00176 for further studies

As mentioned earlier, we primarily focused on LINC007176 to understand the possible regulation of lncRNAs by Δ40p53 and p53. To directly investigate the regulation, the expression of LINC00176 was examined in three cancer cell lines: H1299 cells (null p53 or Δ40p53), HCT116+/+ (endogenous expression of WT p53) and HCT116−/− cells (endogenous expression of Δ40p53). LINC00176 levels were first validated in H1299 cells after overexpressing p53, Δ40p53, or both p53 and Δ40p53 (14A condition). It was upregulated under all conditions compared to control cells **(Figure 1A, B)**. However, the fold change in LINC00176 levels was higher in cells expressing Δ40p53 alone compared to p53 alone or the combination of the two proteins. Since it would be more physiologically relevant to examine LINC00176 in a cell line with endogenous expression of p53 and Δ40p53, its levels were measured in HCT116+/+ and HCT116−/− cells. Likewise, we observed the levels of LINC00176 to be higher in HCT116−/− compared to HCT116+/+ cells **(Figure 1C, D)**. To confirm the ability of Δ40p53 or p53 in regulating LINC00176, an siRNA targeting the 3′-UTR of *TP53* was transfected into HCT116+/+ and HCT116−/− cells **(Figure 1F, H)**. The levels of LINC00176 decreased in both cell lines after siRNA transfection **(Figure 1E, G)**. However, the decrease was not noteworthy because Δ40p53 or p53 are not the only regulators of LINC00176. Nevertheless, silencing Δ40p53 in HCT116−/− cells **(Figure 1G)** led to a more significant reduction in LINC00176 levels than silencing p53 in HCT116+/+ cells **(Figure 1E)**. To further examine whether both isoforms could modulate the levels of LINC00176, p53 was overexpressed in HCT116−/− cells **(Figure 1J)**, which reduced LINC00176 levels compared to the control condition expressing only endogenous Δ40p53 **(Figure 1I)**.

**Figure 1.**
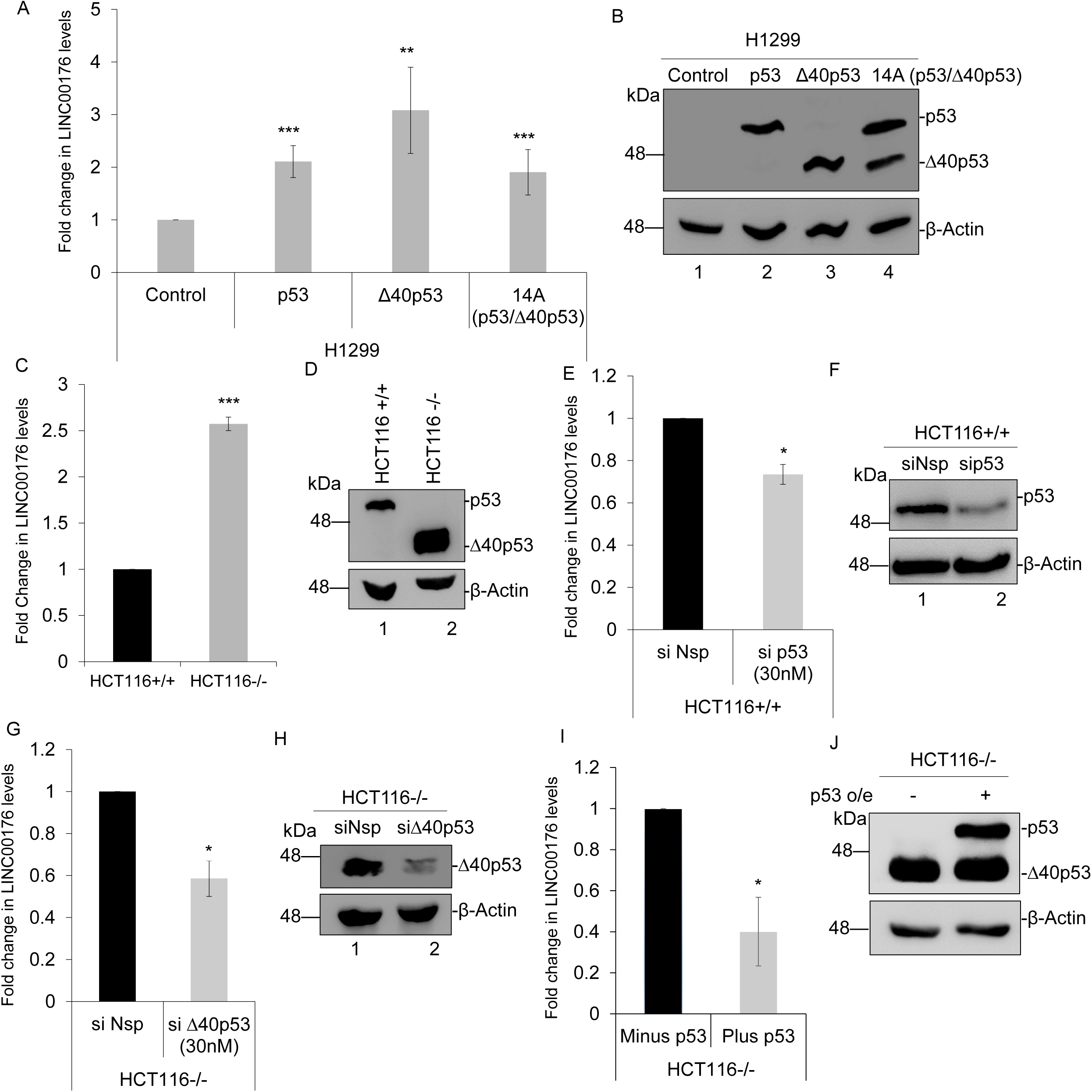
Selection and validation of LINC00176 for further studies. (A) Quantitative PCR for validation of LINC00176 in H1299 cells expressing control, p53 only, Δ40p53 only and 14A construct. (B) Western blot analysis of cell extracts from H1299 cells expressing control, p53 only, Δ40p53 only and 14A construct, probed with CM1 after 48 h. (C) Quantitative PCR for validations of LINC00176 in HCT116+/+ and HCT116−/−cells. (D) Western blot analysis of cell extracts from HCT116+/+ and HCT116−/−cells, probed with CM1. (E) Quantitative PCR of LINC00176 in HCT116+/+ cells transfected with si p53 (30nM) and non-specific si (si Nsp). (F) Western blot analysis of cell extracts from HCT116+/+ cells transfected with either si p53 (30nM) and non-specific si (si Nsp), probed with CM1. (G) Quantitative PCR of LINC00176 in HCT116−/− cells transfected with si Δ40p53 (30nM) and non-specific si (si Nsp). (H) Western blot analysis of cell extracts from HCT116−/− cells transfected with siΔ40p53 (30nM) and non-specific si (si Nsp), probed with CM1. (I) Quantitative PCR of LINC00176 in HCT116−/− cells transfected with control and p53 only. (J) Western blot analysis of cell extracts from HCT116−/− cells transfected with control and p53 only, probed with CM1. The criterion for statistical significance was *p≤*0.05 (*) or *p≤*0.01 (**) or p*≤* 0.001(***)

In H1299 cells, LINC00176 levels increase maximally with Δ40p53 overexpression (3^rd^ bar), while with overexpression of construct 14A (both p53 and Δ40p53), LINC00176 levels are lower than Δ40p53-alone (4^th^ bar), suggesting that in the presence of Δ40p53, p53 negatively affects Δ40p53-mediated LINC expression (4^th^ bar vs. 3^rd^ bar). In HCT116−/− cells, overexpressing p53 reduces LINC00176 in a scenario that resembles the 4^th^ bar vs. the 3^rd^ bar in Figure 1A. The results suggested that both p53 translational isoforms regulate LINC00176, but the lncRNA levels differ concerning isoform abundance.

### Regulation of LINC00176 under different stress conditions

An IRES-mediated mechanism translates p53 and Δ40p53 under stress conditions, such as DNA damage, ER stress, and glucose starvation (Grover, Ray et al. 2008, Sharathchandra, Lal et al. 2012, Khan, Katoch et al. 2015). Cells were treated with doxorubicin and thapsigargin for 16 h to induce DNA damage and ER stress, respectively, and cultured in glucose-deprived DMEM for 8 h to induce glucose starvation **(Figure 2A)**. In HCT116−/− cells, Δ40p53 levels increased under all stress conditions **(Figure 2C, E, G)**. LINC00176 levels also increased, with the highest increase observed in cells treated with thapsigargin **(Figure 2B, D, F)**. In HCT116+/+ cells, although p53 levels increased under all the stress conditions, LINC00176 levels were inconsistent and could not be correlated with p53 levels. It was slightly upregulated in the doxorubicin-treated and glucose-deprived cells; however, there was no significant change in cells treated with thapsigargin **(Figure S1 A-F)**. Moreover, it is unclear whether p53 or Δ40p53 was responsible for the increase in LINC00176 in HCT116+/+ cells, as clearly visible in the case of Doxorubicin induction (**Figure S1 B**) the levels of both the proteins increased. However, in line with our earlier observation of the dominant regulation of LINC00176 by Δ40p53, the results in HCT116−/− cells were more significant and consistent.

**Figure 2.**
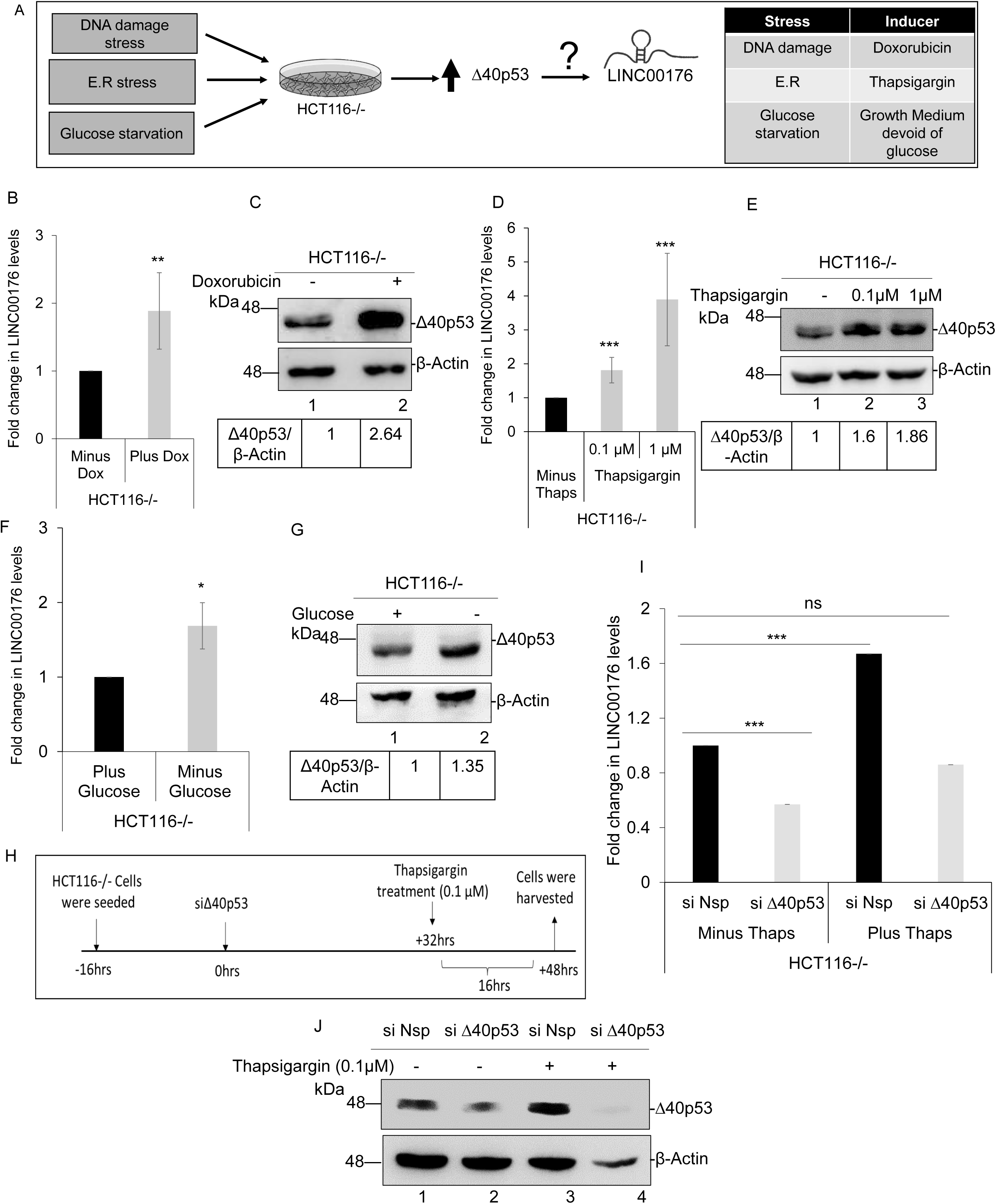
Regulation of LINC00176 under different stress conditions. (A) Schematic of different stress induction in HCT116−/− cells. (B) Quantitative PCR of LINC00176 in HCT116−/− treated with doxorubicin (2µM) for 16 h. (C) Western blot analysis of cell extracts from HCT116−/− treated with doxorubicin (2µM) for 16 h, probed with CM1. (D) Quantitative PCR of LINC00176 in HCT116−/− treated with thapsigargin (0.1µM and 1µM) for 16 h. (E) Western blot analysis of cell extracts from HCT116−/− treated with thapsigargin (0.1µM and 1µM) for 16 h, probed with CM1. (F) Quantitative PCR of LINC00176 in HCT116−/− deprived of Glucose in medium for 8 h. (G) Western blot analysis of cell extracts from HCT116−/− deprived of Glucose in medium for 8 h, probed with CM1. (H) Schematic of experiment in HCT116−/− transfected with siΔ40p53 followed by Thapsigargin (0.1µM) treatment. (I) Quantitative PCR of LINC00176 in HCT116−/− transfected with siΔ40p53 followed by Thapsigargin (0.1µM) treatment. (J) Western blot analysis of cell extracts from HCT116−/− transfected with siΔ40p53 followed by Thapsigargin (0.1µM) treatment, probed with CM1. The criterion for statistical significance was *p≤*0.05 (*) or *p≤*0.01 (**) or p*≤* 0.001(***)

Next, we wanted to confirm whether Δ40p53 alone was involved in regulating LINC00176 levels under stress or whether another factor could also regulate LINC00176. ER stress was selected as it maximally induced Δ40p53 in this model, and we observed significant upregulation of LINC00176 in cells treated with thapsigargin **(Figure 2D)**. Δ40p53 was partially silenced in HCT116−/− cells, followed by induction of ER stress **(Figure 2H)**. Compared to control cells that were untreated and not transfected with siΔ40p53, there was no change in LINC00176 levels in the cells transfected with siΔ40p53 and treated with thapsigargin **(Figure 2I-J)**. This observation indicated that in the absence of Δ40p53, LINC00176 levels are not induced in the background of ER stress, confirming Δ40p53 to be a significant regulator of LINC00176. These results suggest that the expression of LINC00176 is closely linked to Δ40p53 levels. However, the possibility that other factors might also be involved in regulating LINC00176 expression cannot be ruled out. We have also confirmed the ER stress induction by checking the ER chaperone Bip levels (Lee 2005), which regulates the stress (**Figure S1 G**).

### Mechanism of LINC00176 regulation by Δ40p53 or p53

We then investigated whether LINC00176 was directly regulated by Δ40p53 or p53 **(Figure 3A)**. Using PROMO-ALGGEN, we found several Δ40p53 or p53 binding sites in the *LINC00176* gene, suggesting that Δ40p53 or p53 may regulate LINC00176 at the transcriptional level. An earlier study of LINC00176 revealed three putative promoter regions in different cell types (Zhu, Li et al. 2016, Tran, Kessler et al. 2018). ChIP experiments were used to examine the association of Δ40p53 or p53 with these three probable promoter regions: one located in the proximal region of LINC00176 exon 1 (62665399–62666196) and two located in the proximal region of E2 (62667411–62667655 and 62667984–62668233). HCT116−/−and HCT116 +/+ cross-linked lysates were immunoprecipitated with the 1801 antibody, which can detect the Δ40p53 and p53 proteins **(Figure S2A, B)**. The results showed an association between Δ40p53 and *LINC00176* promoters in HCT116−/− cells **(Figure 3B)**. However, no such association was observed for p53 in HCT116+/+ cells **(Figure 3C)**. Since we observed a decrease in LINC00176 levels upon p53 overexpression in HCT116−/− cells **(Figure 1I)**, the same effect of p53 overexpression was examined on the binding of Δ40p53 to *LINC00176* promoters in HCT116−/− cells. Interestingly, results show that the binding of Δ40p53 to all three *LINC00176* promoters decreases upon p53 overexpression in HCT116−/− cells **(Figure 3D, E)**. This observation aligns with the literature showing that p53 and Δ40p53 regulate each other’s activity by forming hetero-oligomers (4). Given the multiple regulators obtained for LINC00176, we were curious to ascertain its cellular localization. Therefore, cell fractionation was performed in HCT116−/− cells; 60% of the transcripts were found to localize to the nucleus, while 40% localized to the cytoplasm **(Figure S2 C)**.

**Figure 3.**
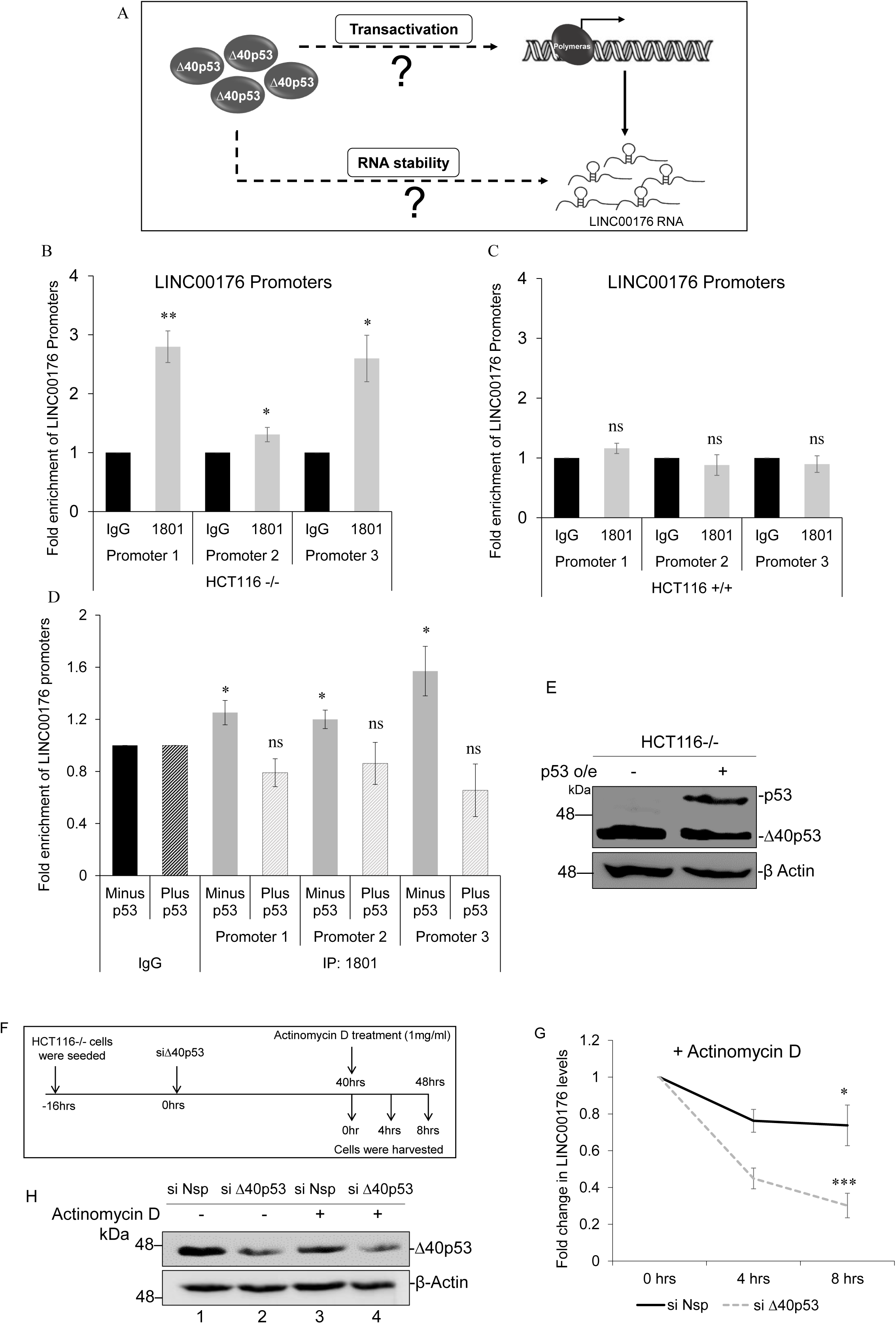
Mechanistic basis of LINC00176 regulation by Δ40p53 or p53. (A) Schematic of possible routes of LINC00176 regulation. (B) Quantitative PCR of LINC00176 promoters in HCT116−/− cells processed for ChIP Assay 48 h post seeding. Δ40p53 was pulled down with 1801 Ab (Santa Cruz) and associated LINC00176 promoters were probed using qPCR. (C) Quantitative PCR of LINC00176 promoters in HCT116+/+ cells processed for ChIP Assay 48 h post seeding. p53 was pulled down with 1801 Ab (Santa Cruz) and associated LINC00176 promoters were probed using qPCR. (D) Quantitative PCR of LINC00176 promoters in HCT116−/− cells transfected with p53 only and control, which has been processed for ChIP Assay 48 h post transfection. Δ40p53 or p53 was pulled down with 1801 Ab (Santa Cruz) and associated LINC00176 promoters were probed using qPCR. (E) Western blot analysis of cell extracts from HCT116−/− transfected with p53 only and control, probed with CM1. (F) Schematic of experiment to determine Δ40p53 mediated regulation on LINC00176 RNA stability. (G) Quantitative PCR of LINC00176 in HCT116−/− transfected with siΔ40p53 followed by Actinomycin D treatment for 0 h, 4 h and 8 h. (H) Western blot analysis of cell extracts from HCT116−/− transfected with siΔ40p53 followed by Actinomycin D treatment, probed with CM1. The criterion for statistical significance was *p≤*0.05 (*) or *p≤*0.01 (**) or p*≤* 0.001(***)

### Deciphering the mechanism of action of LINC00176

Thus far, the results obtained in this study indicate that Δ40p53 can regulate lncRNAs in addition to previously reported miRNAs (Katoch, Tripathi et al. 2021). Therefore, it is crucial to understand how LINC00176 functions in cells and what could be its mechanism of action **(Figure 4A)**. LncRNAs have recently gained attention as competing endogenous RNAs (ceRNAs) known to interact with miRNAs, acting as a decoy or promoting their degradation, thereby liberating mRNAs that directly impact various cellular processes in carcinogenesis (cell proliferation, cell growth, metastasis, etc.) (Yoon, Abdelmohsen et al. 2014, Fan, Sheng et al. 2020, Cui, Pu et al. 2021, Guo, Qian et al. 2021). Therefore, miRNA-interacting partners of LINC00176 were predicted using Starbase V.2.0 database **(Figure 4B)**, and their levels were measured in HCT116+/+ and HCT116−/− cells **(Figure 4C)**. The expression levels of these miRNAs (miR-761, miR-184, miR-503-5p, miR-15b-5p, miR-138, and miR-9-5p) varied. Since most had lower expression in HCT116−/− cells than in HCT116+/+ cells, they could be targets of LINC00176, as LINC00176 levels were very high in HCT116−/− cells. In particular, miR-9-5p, a known target of LINC00176 (Tran, Kessler et al. 2018), was downregulated in HCT116−/− cells compared with its expression in HCT116+/+ cells. To assess the direct effect of LINC00176 on these miRNAs, LINC00176 was silenced in HCT116−/− cells and the corresponding changes in miRNA levels were measured. Upon silencing LINC00176, the levels of LINC00176 fell by 60% compared with those in the control cells **(Figure 4D)**. The miRNA levels increased from 30% to 80% **(Figure 4E)**. To investigate whether LINC00176 sequesters target miRNAs, a tagged RNA affinity pull-down assay was performed **(Figure 4G)**. miRNA levels were quantified in the pull-down fractions of MS2-control and MS2-LINC00176 and normalized to their respective input levels. Furthermore, the normalized miRNA levels in MS2-LINC00176 were compared with those in MS2-Control to calculate the fold enrichment. Results revealed that miR-184, miR-138, miR-503, miR-15b-5p, and miR-9-5p were significantly enriched in the pull-down fraction, indicating that they were direct targets of LINC00176 **(Figure 4F)**. However, miR-761 was not enriched in the pull-down fraction.

**Figure 4.**
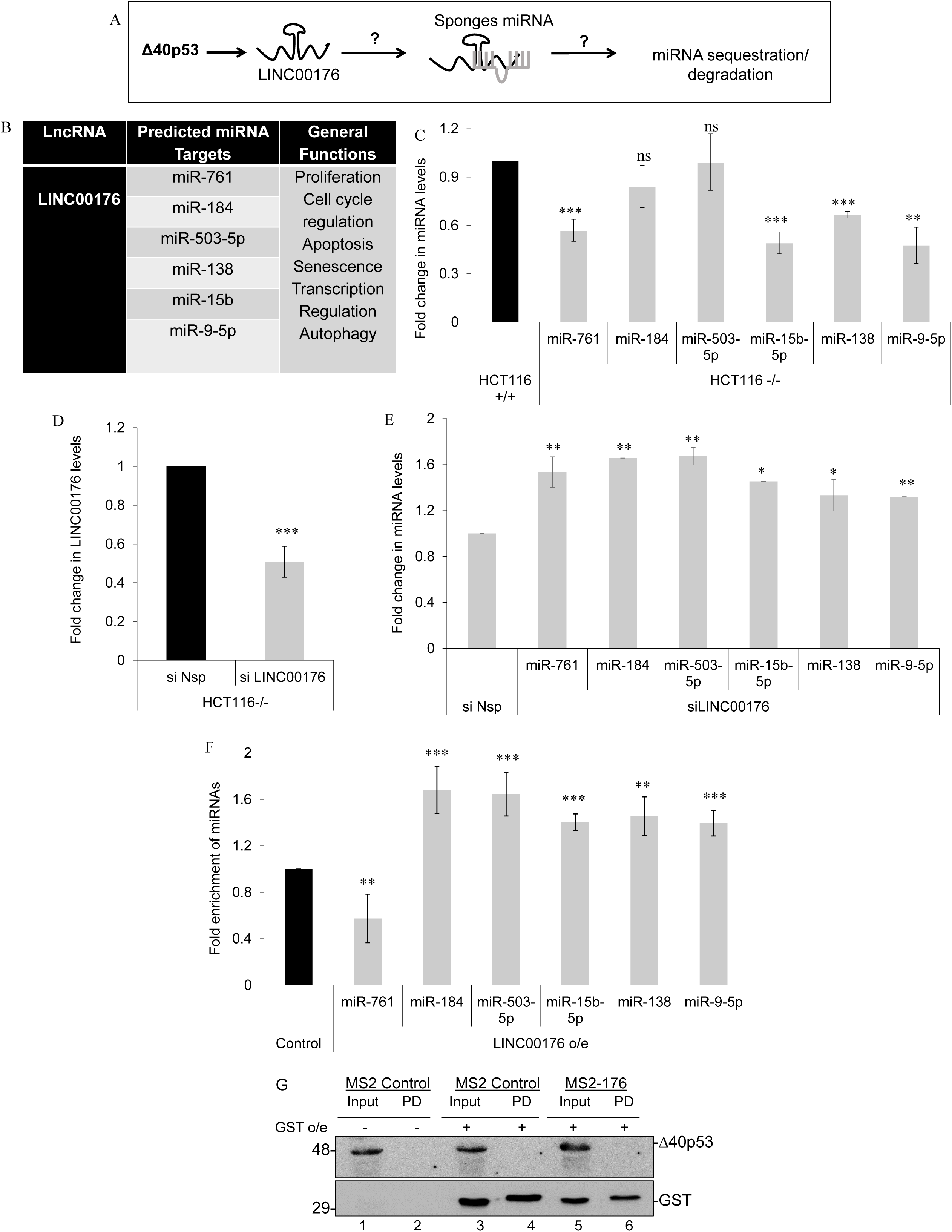
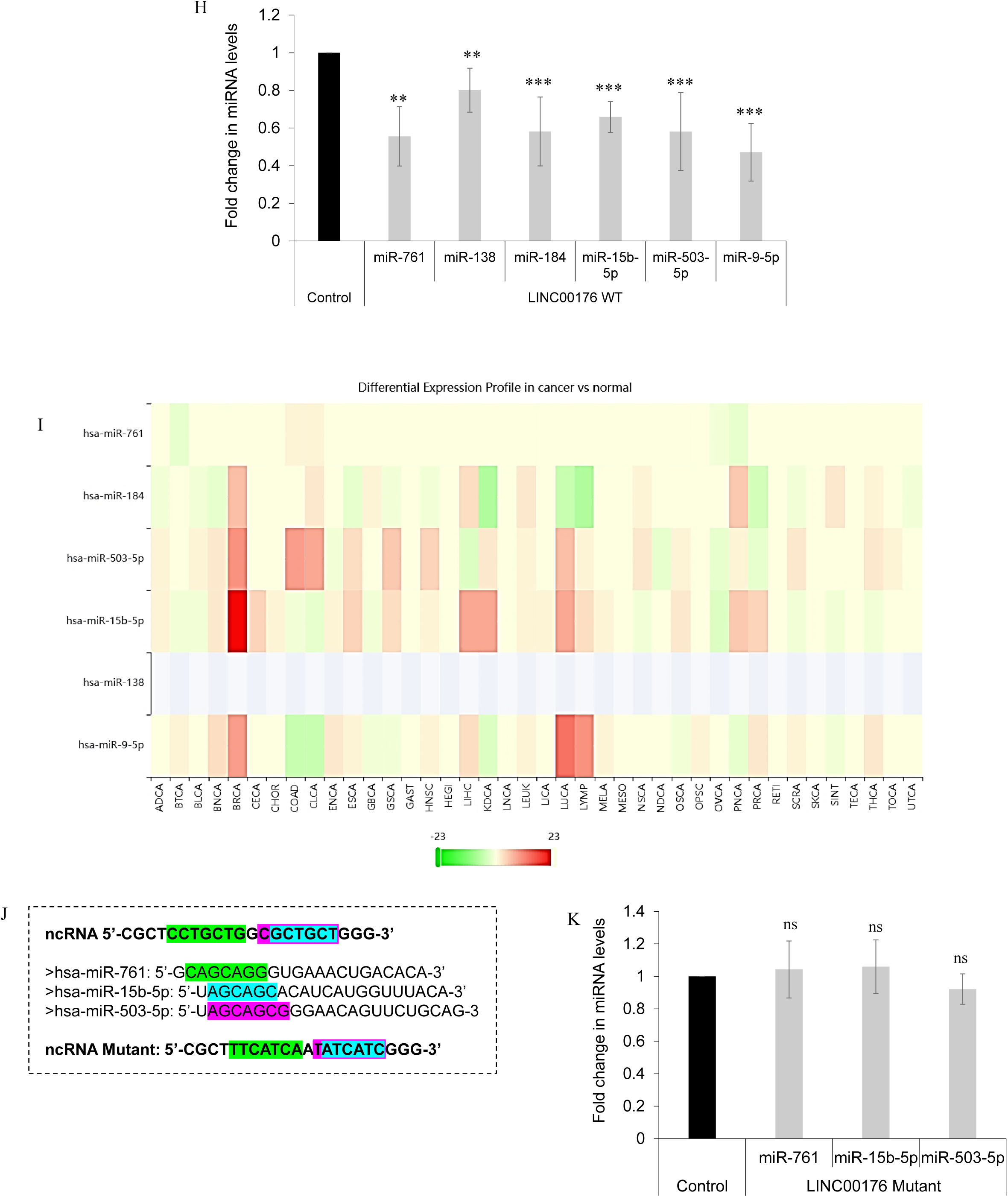
Deciphering the mechanism of action of LINC00176. (A) Schematic for the probable downstream regulation by LINC00176. (B) Table for the selected miRNA targets of LINC00176 obtained from Starbase V2.0, as mentioned in methods section. (C) Quantitative PCR of selected miRNAs in HCT116+/+ and HCT116−/−cells. (D) Quantitative PCR of LINC00176 in HCT116−/− cells transfected with siRNA (30nM) directed to LINC00176 and non-specific si (si Nsp). (E) Quantitative PCR of miRNAs in HCT116−/− cells transfected with siRNA (30nM) directed to LINC00176 and non-specific si (si Nsp). (F) Quantitative PCR of selected miRNAs in GST pulldown lysates of HCT116−/−cells (co-transfected with LINC00176 overexpression construct and pMS2-GST construct) normalised to Input values. (G) Western blot analysis of cell extracts from Input and Pulldown fraction of HCT116−/− cells (co-transfected with LINC00176 overexpression construct and pMS2-GST construct), probed with CM1 and GST Antibody. (H) Quantitative PCR of miRNAs in HCT116−/− cells transfected with LINC00176 overexpression construct. (I) Differential levels of the selected miRNAs in cancer obtained from dbDEMC 3.0 database. (J) Schematic of the mutations generated for miRNA binding sites on LINC00176. Green colour, blue colour and pink colour signifies overlapping binding sites for miR-761, miR-15b-5p and miR-503-5p respectively. (K) Quantitative PCR of miR-761, miR-15b-5p and miR-503-5p in HCT116−/− cells transfected with LINC00176 mutant construct. The criterion for statistical significance was *p≤*0.05 (*) or *p≤*0.01 (**) or p*≤* 0.001(***)

We also overexpressed LINC00176 and directly checked miRNA levels. Upon LINC00176 overexpression, the miRNAs were downregulated by 40–50%, implying that miRNAs sequestered by LINC00176 could also be degraded in the complex **(Figure 4H)**. Additionally, the specificity of the binding of miRNAs to LINC00176 was examined. Three miRNAs (miR-503-5p, miR-15b-5p, and miR-761) were selected based on their known functions in cancer **(Figure 4I)** and because they share overlapping binding sites with LINC00176. To study the specificity, we mutated the entire binding site (comprising sites for all three miRNAs) of LINC00176 **(Figure 4J)**. Upon overexpression of mutant LINC00176, we observed that the miRNA levels were alleviated **(Figure 4K)** compared to the decrease observed in Figure 4H.

LINC00176-mediated differential abundances obtained for miRNAs led us to speculate their miRNA: LINC00176 copy numbers in cells. The copy numbers of LINC00176 versus the miRNAs were analyzed **(Figure S3)**, and we observed that the number of miRNAs present was very high compared with LINC00176 levels **(Figure S3, H)**. Thus, results from Figure 4 E, H and Figure S3 H implied that even though LINC00176 might be lower in cells than miRNAs, it can significantly regulate miRNA abundance. As a consequence of the LINC00176-miRNA regulation, several downstream targets driving critical cellular processes could also be affected.

### Identifying the mRNA players downstream to the LINC00176-mRNA pathway

The direct cellular effect of the LINC00176-miRNA axis was determined by identifying the mRNA targets of the microRNAs. miRTarBase was used to identify unique and commonly validated miRNA targets **(Figure 5A)**. For further investigation, miR-503-5p and miR-15b-5p were selected, which have variable abundance in several cancer conditions **(Figure 4I)**. miR-9-5p (as a positive control) was also used as it is an established target of LINC00176 **(Tran, Kessler et al. 2018)** and shares several mRNA targets with the other two miRNAs. To assess the direct effect of LINC00176 on these miRNA targets, LINC00176 was silenced in HCT116−/− cells and the corresponding changes in mRNA levels were measured. Upon silencing, mRNA levels varied differentially, with a maximum decrease in the levels of Bcl2 mRNA **(Figure 5B)**. Moreover, mRNA levels were examined against the backdrop of 40p53, the main regulator of the axis. After silencing Δ40p53 in HCT116−/−, all the target levels changed differently, with a consistent decrease in CCNE1, SIRT1, and Bcl2 levels **(Figure 5 C-D)**, corroborating the results obtained in the siLINC00176 background.

**Figure 5.**
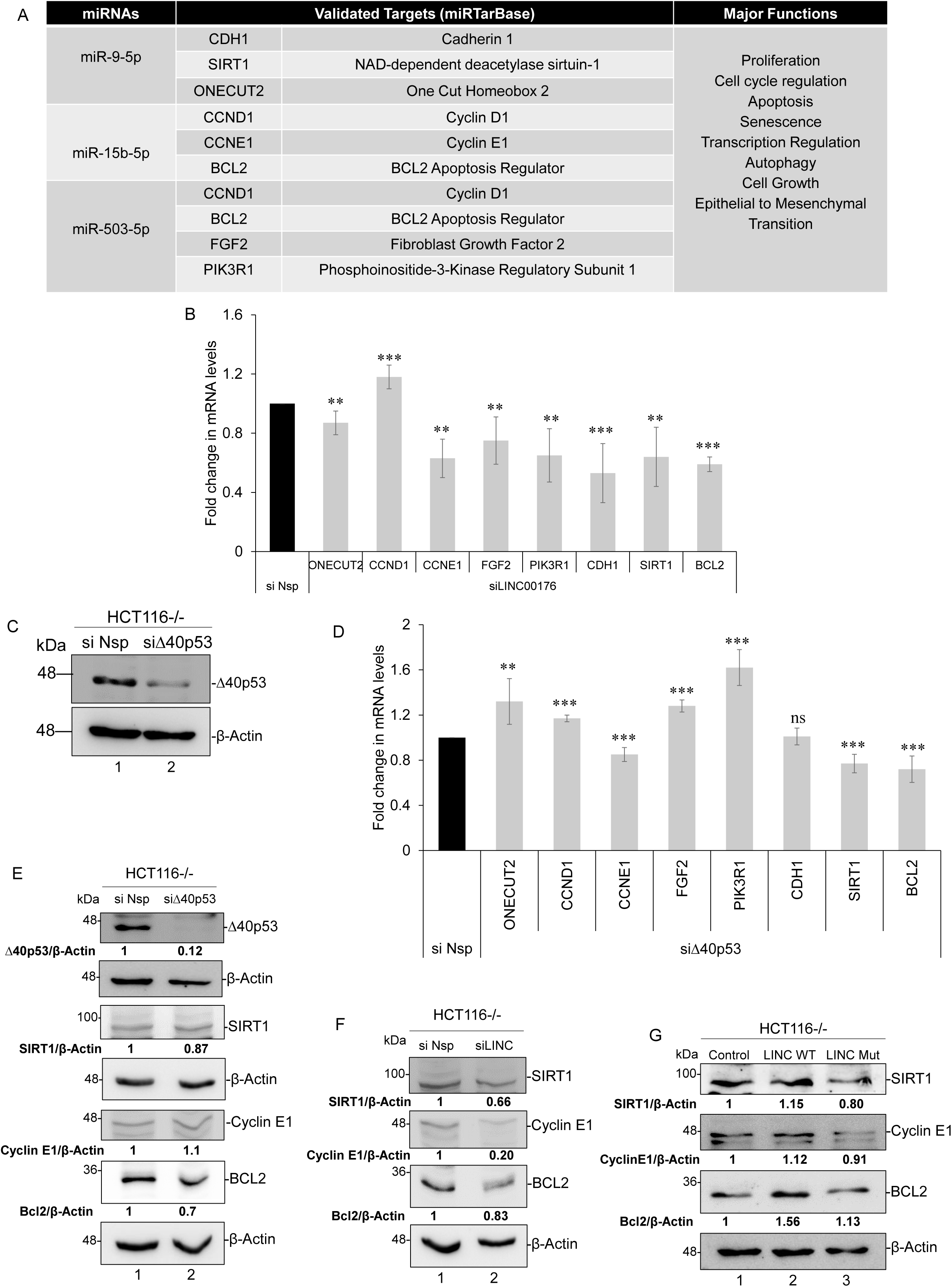
Identifying the potential mRNA targets of miRNAs. (A) Table for the selected mRNA targets of miRNAs obtained from miRTarBase, as mentioned in methods section. (B) Quantitative PCR of selected mRNAs in HCT116−/− cells transfected with siRNA (30nM) directed to LINC00176 and non-specific si (si Nsp). (C) Western blot analysis of cell extracts from HCT116−/− cells transfected with siΔ40p53 (30nM) and non-specific si (si Nsp). (D) Quantitative PCR of selected mRNAs in HCT116−/− cells transfected with siΔ40p53 (30nM) and non-specific si (si Nsp). (E) Western blot analysis of cell extracts from HCT116−/−cells transfected with siΔ40p53, probed with SIRT1, Cyclin E1, BCL2 and 1801 Antibody. (F) Western blot analysis of cell extracts from HCT116−/−cells transfected with siLINC00176, probed with SIRT1, Cyclin E1, BCL2 and 1801 Antibody. (G) Western blot analysis of cell extracts from HCT116−/−cells transfected with Control, WT LINC00176 and Mutant LINC00176 probed with SIRT1, Cyclin E1, BCL2 and 1801 Antibody. The criterion for statistical significance was *p≤*0.05 (*) or *p≤*0.01 (**) or p*≤* 0.001(***).

Since proteins are the ultimate regulators of cellular pathways, CCNE1, SIRT1, and Bcl2 protein levels were also examined upon silencing Δ40p53 and LINC00176. We observed a consistent decrease in BCL2 and SIRT1 protein levels under both conditions, whereas Cyclin E1 levels only decreased under siLINC00176 **(Figure 5 E-F)**. Moreover, the protein levels were also checked in the background of WT LINC00176 and mutant LINC00176 (as explained above). SIRT1, CCNE1, and Bcl2 levels increased upon WT LINC00176 overexpression, whereas they were restored to normal levels upon overexpression of mutant LINC00176 **(Figure 5G)**.

As observed earlier, overexpression of WT LINC00176 reduced the levels of all miRNAs **(Figure 4H);** thus, it upregulated the targets of miR-15b-5p and miR-503-5p (also shared for miR-9-5p) **(Figure 5G, lane 2)**. Mutant LINC00176 did not affect the levels of miR-15b-5p and miR-503-5p **(Figure 4K)**; thus, the levels of the protein targets (SIRT1, Bcl2, and CCNE1) were restored to normal levels **(Figure 5 G, lane 3)**. These differential abundances of the proteins under different backgrounds of LINC00176 helped unravel the distant players of the Δ 40p53-LINC00176-miRNA pathway, which might directly affect the cellular processes.

### The impact of the Δ40p53-LINC00176 axis on cellular processes

LncRNAs regulate a wide variety of cellular processes (Statello, Guo et al. 2021). The ceRNA network involving lncRNA/miRNA/mRNA synergistically regulates tumor hallmark processes across pan cancers (Zhang, Xu et al. 2016). A lncRNA can have multiple networks in cancer (Wang, Cho et al. 2019); therefore, lncRNAs being the master regulator, it is crucial to know their ultimate impact on cells (primarily on cell growth, expression of EMT markers, viability etc.). To understand the direct effect of LINC00176 on cellular behavior, LINC00176 was over-expressed or knocked down in HCT116−/− cells **(Figure 6 A-C)**. First, we examined the effect of LINC00176 on EMT marker expression. LINC00176 overexpression increased the levels of E-cadherin (epithelial marker) and decreased the levels of Slug and vimentin (mesenchymal markers) **(Figure 6 D, E)**. By contrast, knocked-down LINC00176 decreased the mRNA levels of E-cadherin but increased those of vimentin and Slug **(Figure 6 F);** however, the same trend was not observed for the protein levels of E-cadherin and Slug **(Figure 6 G)**. We further analyzed cell viability after LINC00176 overexpression or knockdown and observed that LINC00176 overexpression decreased cell viability **(Figure 6 H)**, whereas LINC00176 knockdown increased cell viability **(Figure 6 I)**. However, these effects on cell viability may not be very prominent because LINC00176 might contribute to only a tiny percentage of the vast majority of cellular processes. LINC00176 knockdown also increased proliferation in the BrdU assay (**Figure 6 J)**. Together, these results indicate that LINC00176 promotes the epithelial phenotype and suppresses the mesenchymal phenotype, cell growth, viability, and proliferation, all of which suggest that this lncRNA contributes to tumor suppression in cells.

**Figure 6:**
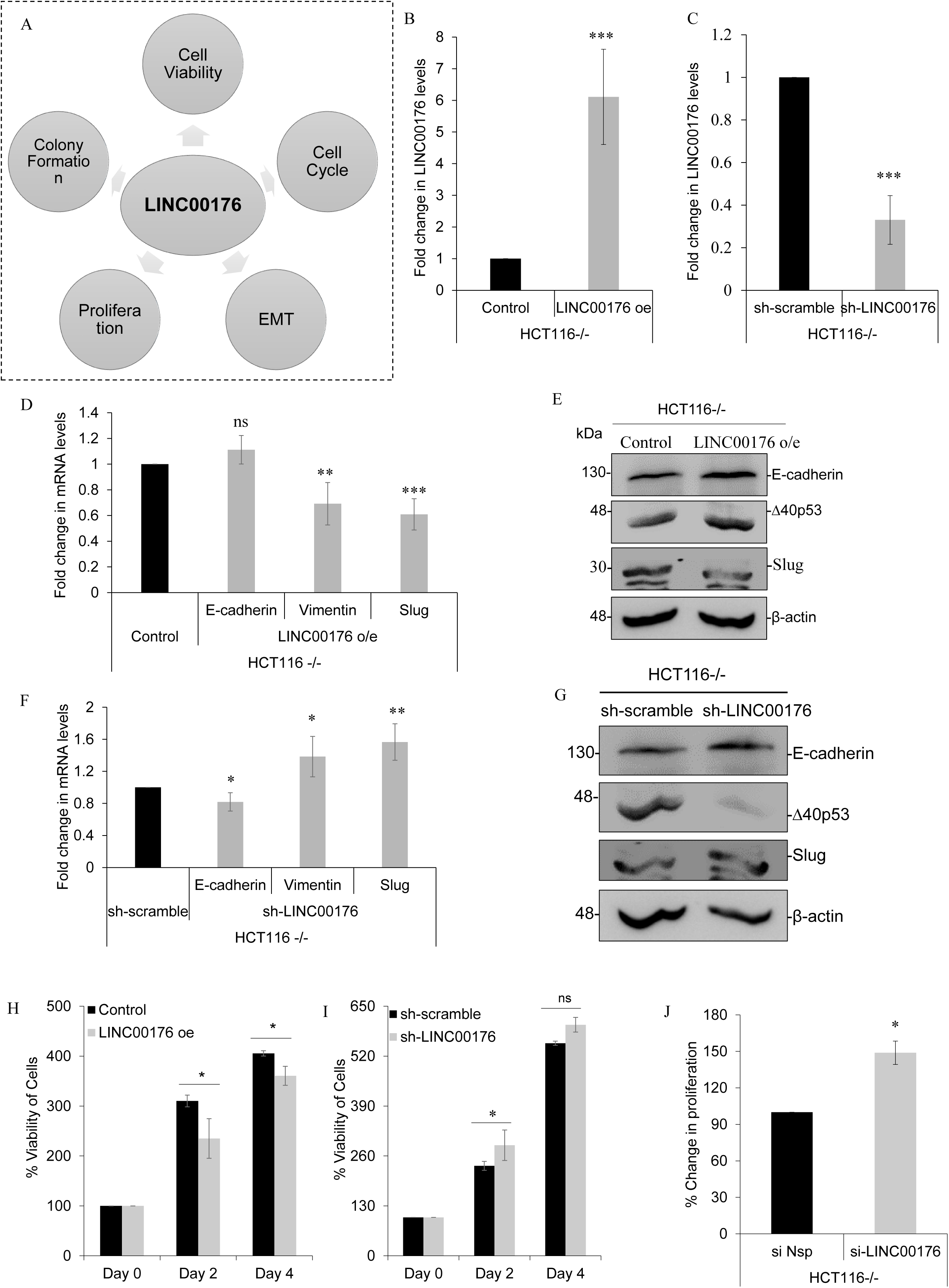
The impact of Δ40p53-LINC00176 axis on cellular processes. (A) Schematic of the different processes that has been checked for any probable effect by LINC00176 over-expression or partial silencing. (B) Quantitative PCR of LINC00176 in HCT116−/− cells transfected with LINC00176 overexpression construct. (C) Quantitative PCR of LINC00176 in HCT116−/− cells transfected with LINC00176 shRNA construct. (D) Quantitative PCR of EMT markers (E-cadherin, Vimentin, Slug) in HCT116−/− cells transfected with LINC00176 overexpression construct. (E) Western blot analysis of cell extracts from HCT116−/−cells transfected with LINC00176 overexpression construct, probed with E-cadherin Antibody, Slug Antibody and CM1. (F) Quantitative PCR of EMT markers (E-cadherin, Vimentin, Slug) in HCT116−/− cells transfected with LINC00176 shRNA construct. (G) Western blot analysis of cell extracts from HCT116−/−cells transfected with LINC00176 shRNA construct, probed with E-cadherin Antibody, Slug Antibody and CM1. (H) HCT116−/− cells were transfected with LINC00176 overexpression construct, at 36 h post transfection cells were re-seeded on 96 well plate for MTT Assay. MTT incorporation was calculated at Day 0, Day 2 and Day 4 post seeding. (I) HCT116−/− cells were transfected with LINC00176 shRNA construct, at 36 h post transfection cells were re-seeded on 96 well plate for MTT Assay. MTT incorporation was calculated at Day 0, Day 2 and Day 4 post seeding. (J) HCT116−/− cells were transfected with 30nM si-LINC00176, 36 h post transfection cells were re-seeded for BrdU assay. After 12 h of seeding BrdU was added to the cells and incubated for 12 h. Following which, percentage of cell proliferation was checked by calculating BrdU incorporation. The criterion for statistical significance was *p≤*0.05 (*) or *p≤*0.01 (**) or p*≤* 0.001(***).

## DISCUSSION

In cancer, lncRNAs are vital players in the p53-dependent transcriptional pathway, wherein lncRNAs and p53 can regulate each other (Lin, Hou et al. 2019). Therefore, it is vital to recognize novel lncRNAs regulated through the p53 pathway to identify critical cancer signaling pathways. *TP53* is the most frequently mutated gene in human cancer (Goldstein, Marcel et al. 2011); however, these mutations are not found in all cancers, indicating that other factors could interfere with WT p53 function (Zhang, Groen et al. 2022). These factors have complicated our understanding of p53 signaling and include more recently discovered p53 isoforms (Vieler and Sanyal 2018). Owing to the lack of detailed functional characterization, little information is available regarding p53 isoforms. However, one isoform that has recently gained attention in cancer research is Δ40p53, the only translational isoform of p53 (Steffens Reinhardt, Zhang et al. 2020). Although a few reports have shown the transactivation capacity of Δ40p53, there are no studies on the regulation of lncRNAs by Δ40p53. We selected LINC00176 for our current study because of its variable regulation by different transcription factors and differential expression in cancer.

Significant changes in the levels of LINC00176 were observed in H1299 cells expressing p53, Δ40p53, and both isoforms **(Figure 1A)**. However, the maximum upregulation was observed in cells expressing only Δ40p53 **(Figure 1A, 3rd bar)**. The upregulation of LINC00176 by Δ40p53 was also observed in other cell lines expressing only Δ40p53 (HCT116−/−) as compared to cells expressing both isoforms (HCT116+/+) **(Figure 1C)**. Furthermore, LINC00176 was upregulated under different stress conditions, and its expression correlated with the abundance of Δ40p53 **(Figure 2)**. The regulation by Δ40p53 could be at the transcriptional level or post-transcriptional; therefore, we investigated any direct binding between the molecules.

Δ40p53 was previously thought to be devoid of transactivation capacity because it lacks TAD1 (Courtois, Verhaegh et al. 2002); however, subsequent studies reported that it could regulate some p53 target genes (Yin, Stephen et al. 2002, Ohki, Kawase et al. 2007). It has also been reported that Δ40p53 can induce cell death independent of p53 through a specific subset of genes (Fas, Dr5, Api1, and Pig3) (Phang, Othman et al. 2015). More recently, a study suggested an FLp53-independent, pro-tumoral role for Δ40p53 because of its ability to transactivate the transcription of antiapoptotic proteins, such as netrin-1 (Sun, Manceau et al. 2021). Therefore, we examined the association of Δ40p53 and p53 with LINC00176. We observed the binding of Δ40p53 to *LINC00176* promoters **(Figure 3B)**; surprisingly, p53 did not bind to *LINC00176* promoters **(Figure 3C)**. Overexpression of p53 in HCT116−/− cells decreased the association of Δ40p53 with *LINC00176* promoters, suggesting the formation of hetero-oligomers between p53 and Δ40p53 interfered with the transactivation function of Δ40p53, which is consistent with the results of earlier reports (Ghosh, Stewart et al. 2004). Besides the role of p53 as a typical transcription factor, there are reports of p53 binding to RNA through a nucleic acid binding domain, which is independent of its core DNA binding domain(Riley and Maher 2007, Riley, Ramirez-Alvarado et al. 2007). Recently it has been found that p53 can bind to the lncRNA MEG3(Zhu, Liu et al. 2015); however, there are no such reports on other p53 isoforms. With the Actinomycin D experiment, we got a clue that Δ40p53 could also regulate the stability of LINC00176 RNA **(Figure 3G)**. The mechanism through which a transcription factor could bind to an RNA is intriguing, which we would like to study it in the future. We see higher enrichment of the promoters in the background of Δ40p53 alone compared to when both p53 and Δ40p53 are present together **(Figure 3B and C**, respectively). When both isoforms are present, there is a near-complete absence of any promoter-binding or enrichment with Antibody-1801 that can detect either isoform **(Figure 3C)**. The lack of binding of any p53 isoform to the LINC promoter, even when these are present together, is validated when p53 is overexpressed in endogenous-Δ40p53-expressing HCT116−/− **(Figure 3D)**. These collectively lead us to infer that p53 modifies Δ40p53-mediated transactivation of LINC00176 by negatively regulating Δ40p53 activity but without directly binding LINC00176 promoters. Additionally, we find a 60:40 Nuclear: Cytoplasmic ratio of LINC00176 in HCT116−/− **(Figure S2 C)**, which confirms its levels in both compartments, possibly a consequence of its regulation at multiple levels.

The significance of LINC00176 upregulation by Δ40p53 was determined by examining different miRNA targets. We identified different miRNA targets using StarBase V2.0 for LINC00176, from which we selected a few targets (miR-761, miR-138, miR-15b-5p, miR-503-5p, miR-184, and miR-9-5p) based on the number of binding sites and their functions in different cancers. Most of the selected miRNAs are differentially expressed in cancer, owing to their diverse regulatory pathways (Wong, Liu et al. 2008, Yan, Yang et al. 2015, Zhou, Zhang et al. 2016, Li, Wu et al. 2017, Sha, Wang et al. 2017, Li, Li et al. 2018, Dong, Zhang et al. 2019, Wu, Huang et al. 2019, Fei, Shan et al. 2020, Li, Zhang et al. 2020, Liu, Tian et al. 2020, Wu, Liu et al. 2020, Jiang, Wang et al. 2021, Wang, Wu et al. 2021, Wang, Cui et al. 2021, Wen, Feng et al. 2021, Zhu, Lin et al. 2021). Their levels were mainly decreased in HCT116−/− cells compared with those in HCT116+/+ cells, and their expression was negatively correlated with LINC00176 levels **(Figure 4C)**.

Upon further analysis of their levels after knockdown of LINC00176 and TRAP assays, we found that most of these miRNAs (miR-138, miR-15b-5p, miR-503-5p, miR-184, and miR-9-5p) were enriched in the pull-down complex of LINC00176. Although the level of miR-761 was increased upon LINC00176 knockdown, it was not sequestered by LINC00176 **(Figure 4F)**. While knockdown of LINC00176 increased the levels of miRNAs, overexpression of the lncRNA significantly decreased the levels of all miRNAs, indicating a possible mechanism of target RNA-directed miRNA degradation **(Figure 4H)**. We also checked for the specificity of miRNA binding by mutating miR-761, miR-503-5p, and miR-15b-5p miRNA target sites on LINC00176 and observed a rescue in the miRNA levels under mutant LINC00176 expression **(Figure 4K)**. Comparative analysis of the copy number of LINC00176 vs. miRNAs in HCT116−/− cells indicated that the miRNAs were higher in cells than LINC00176. However, when we silenced or overexpressed WT and Mutant LINC00176, the ratio changed, as did the levels of miRNAs, which implies that the amount of LINC00176 increased or decreased is sufficient to modulate miRNA expression.

These results suggest that LINC00176 is an essential ceRNA titrating the levels of multiple miRNAs, thereby attenuating possible miRNA-mRNA interactions. To elucidate the effect of LINC00176 on downstream mRNAs, we validated the mRNA targets of the miRNAs. We used miRTarBase to identify some unique and common validated targets (CDH1, SIRT1, ONECUT2, Bcl2, PIK3R1, FGF2, CCND1, and CCNE1) of the selected miRNAs (miR-9-5p, miR-503-5p, and miR-15b-5p). We knocked down two upstream regulators, Δ40p53 and LINC00176, in HCT116−/− cells and checked the corresponding changes in mRNA and protein levels. Target levels (particularly Bcl2 and SIRT1) decreased upon silencing both upstream regulators and increased upon overexpression of LINC00176, indicating their potential involvement in the Δ40p53/LINC00176/miRNA axis **(Figure 5 B, D, E, F)**. However, a detailed investigation is necessary to correlate the direct effects of individual proteins on downstream processes.

Finally, the direct effects of LINC00176 were examined on cellular behavior through ectopic overexpression or silencing of LINC00176. Based on the different hallmark processes of cancer (Hanahan and Weinberg 2011) we examined EMT marker expression, cell proliferation, and cell viability. LINC00176 overexpression enhanced the epithelial phenotype and decreased cell viability **(Figure 6 E, H)**. LINC00176 knockdown led to an increase in cell viability and proliferation; however, it did not give consistent results with the EMT markers, which also decreased upon LINC00176 knockdown **(Figure 6 I, J and G)**. We speculate that the decrease in the protein levels of EMT markers could be due to a decrease in Δ40p53, which might also directly regulate EMT markers. Therefore, the effect of LINC00176 knockdown is inconclusive. Δ40p53 function is cell context-specific, and Δ40p53 can have differential effects when overexpressed or silenced (Zhang, Groen et al. 2022). This observation strengthens our hypothesis that, since there is a possible feed-forward loop between LINC00176 and Δ40p53, many of the lncRNA effects will depend on the levels of Δ40p53 and, therefore will vary. The mechanistic details of the possible feed-forward loop is an interesting axis we would like to decipher in the the future.

In conclusion, our results indicated that LINC00176 contributes to the tumor-suppressive function of Δ40p53, independent of the p53FL function **(Figure 7: Graphical Abstract)**. However, it would be interesting to determine whether the function of the Δ40p53-LINC00176 axis is universal or cell type-specific. Collectively, this study opens up avenues for mechanistic insights into how Δ40p53 governs cellular processes through lncRNAs.

**Figure 7:**
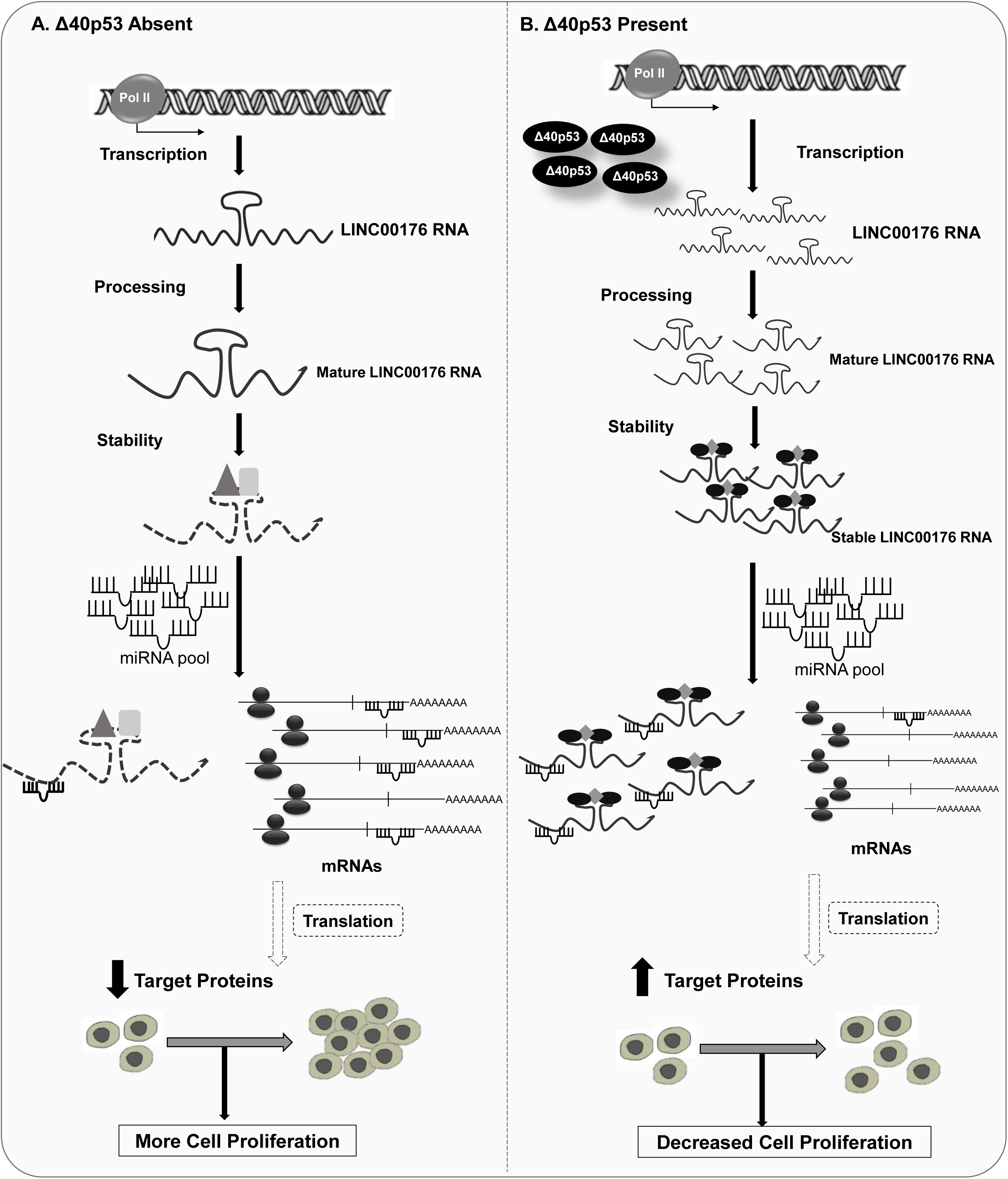
Graphical Abstract: Model depicting the regulation of Δ40p53-LINC00176-miRNA axis. (A) In absence of Δ40p53, LINC00176 levels do not increase which leads to increased pool of free miRNAs which probably binds to several mRNA targets preventing their function, ultimately resulting in increased cell proliferation. (B) However, when Δ40p53 is present it upregulates LINC00176 levels both at the level of transcription and post-transcription, which sequesters oncogenic miRNAs preventing it to bind to their respective mRNA targets, which ultimately results in decreased cell proliferation.

## ACKNOWLEDGEMENTS

We are grateful to Prof. J.C. Bourdon, University of Dundee, UK and Dr. Robin Fahraeus, INSERM, France, for providing us the anti-p53 antibody. We would like to thank Dr. Je-Hyun Yoon and Dr. Myriam Gorospe, National Institutes of Health, USA, for providing us with the TRAP constructs. We are grateful to Dr. T Tamura, Institut fuer Biochemie, Germany, for providing us with LINC00176 overexpression construct and siRNA. We would like to thank Prof. Sathees C. Raghavan for his valuable suggestions. We would also like to acknowledge and sincerely thank Dr. Debjit Khan (Post-Doctoral Research Fellow, Cleveland Clinic, USA) for his invaluable suggestions. We also thank the present and past SD lab-members for helpful discussion of the work. This work is supported by a research grant to SD from the Department of Biotechnology (DBT), Govt. of India. SD also acknowledges J.C. Bose fellowship for research support. This study was also supported by the DBT-IISc partnership program. DST Fund for Improvement of Science and Technology Infrastructure (FIST) level II infrastructure and University Grants Commission Centre of Advanced Studies support is acknowledged. We would like to thank Editage (www.editage.com) for English language editing.

## AUTHOR CONTRIBUTIONS

AP and SD: Conception and design of studies, analysis and interpretation, and article writing. AP and PKG and for performing the experiments.

## DATA AVAILABILITY STATEMENT

Data available within the article or its supplementary materials.

