## Supplementary Figure legends for "The “LINC” in between Δ40p53-miRNA axis in the regulation of cellular homeostasis"

**Figure S1: Regulation of LINC00176 under different stress conditions**

(A) Quantitative PCR of LINC00176 in HCT116+/+ treated with Doxorubicin (2µM) for 16 h. (B) Western blot analysis of cell extracts from HCT116+/+ treated with Doxorubicin (2µM) for 16 h, probed with CM1. (C) Quantitative PCR of LINC00176 in HCT116+/+ treated with Thapsigargin (0.5µM) for 16 h. (D) Western blot analysis of cell extracts from HCT116+/+ treated with Thapsigargin (0.5µM) for 16 h, probed with CM1. (E) Quantitative PCR of LINC00176 in HCT116+/+ deprived of Glucose in medium for 8 h. (F) Western blot analysis of cell extracts from HCT116+/+ deprived of Glucose in medium for 8 h, probed with CM1. (G) Western blot analysis of cell extracts from HCT116-/- transfected with siΔ40p53 followed by Thapsigargin (0.1µM) treatment, probed with Bip Antibody. (P = NS, not significant, *P < 0.05, **P <0.01, *** P < 0.001).

**Figure S2: Mechanistic basis of LINC00176 regulation by Δ40p53/p53**

(A) Western blot analysis of cell extracts from HCT116-/- processed for ChIP Assay 48 h post seeding, probed with 1801 Antibody. (B) Western blot analysis of cell extracts from HCT116+/+ processed for ChIP Assay 48 h post seeding, probed with 1801 Antibody. (C) Relative expression of GAPDH, MALAT1 and LINC00176 transcripts in HCT116-/- cells upon Nuclear-Cytoplasmic fractionation. (Cytoplasmic Control RNA), MALAT1 (Nuclear Control RNA)

**Figure S3. Deciphering the mechanism of action of LINC00176**

(A-G) Standard curves of all miRNAs and LINC00176. X-axes for the graphs represents RNA copy number (log 2 expression of copy number) and Y-axis represents the Ct values obtained for the respective copy number. (H) Table for comparative analysis of the copy numbers obtained for the RNAs with their endogenous expression.
