## Supplementary Figures for "The “LINC” in between Δ40p53-miRNA axis in the regulation of cellular homeostasis"

### Figure: S1

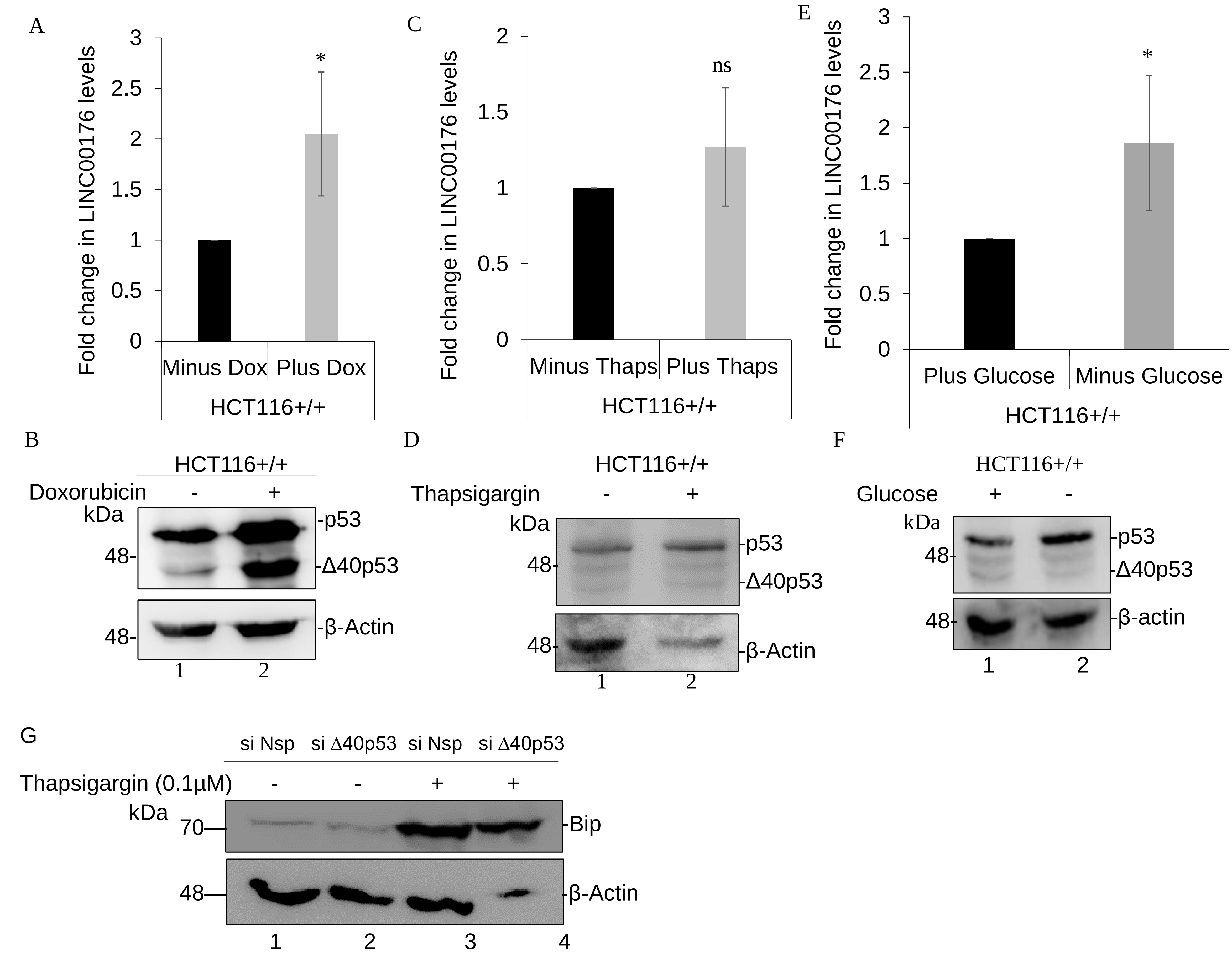


| Bip/β-Actin | 1 | 1.31 | 4.49 | 14.46 |
|---|---|---|---|---|

### Figure: S2

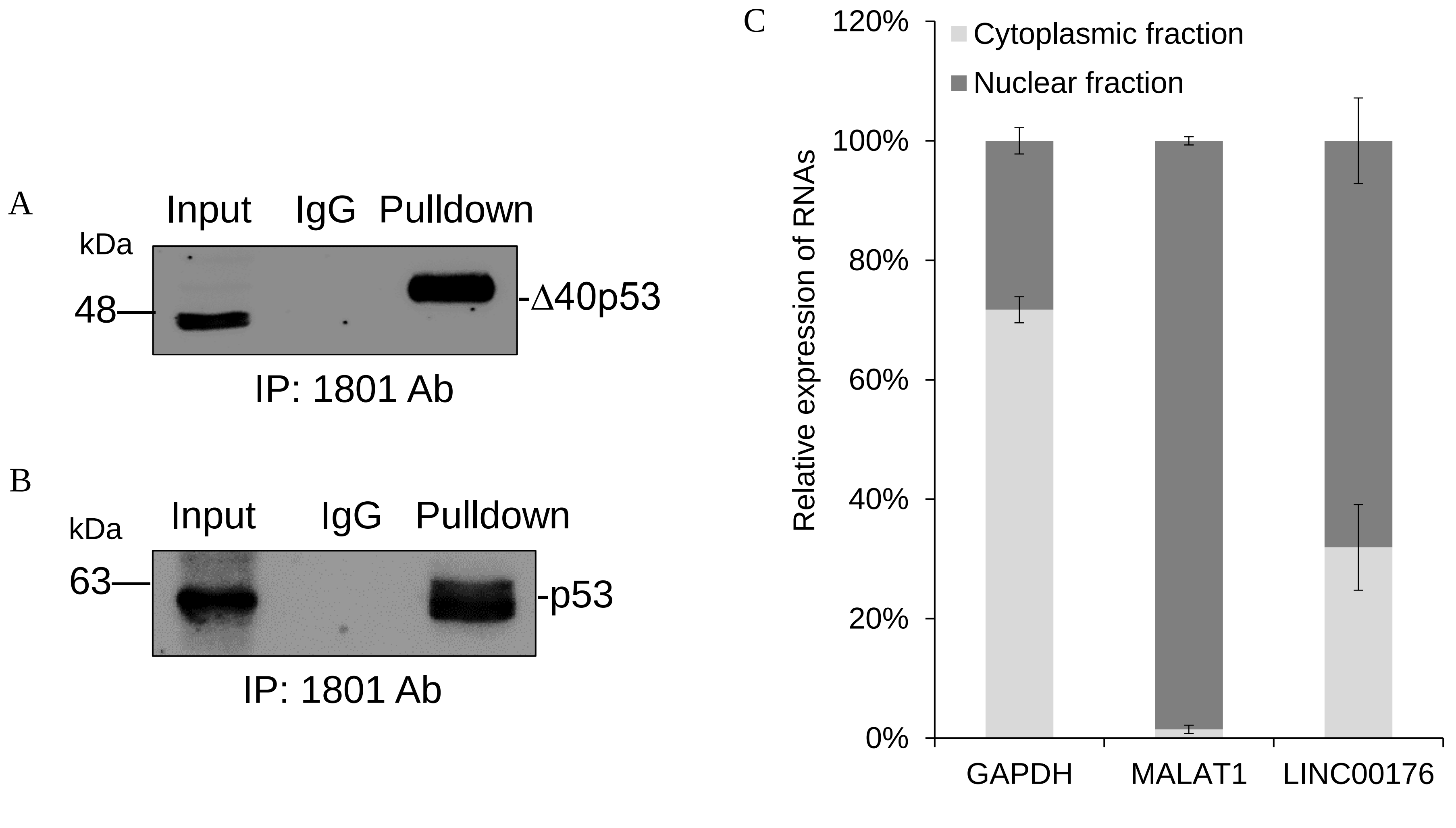

### Figure: S3

A

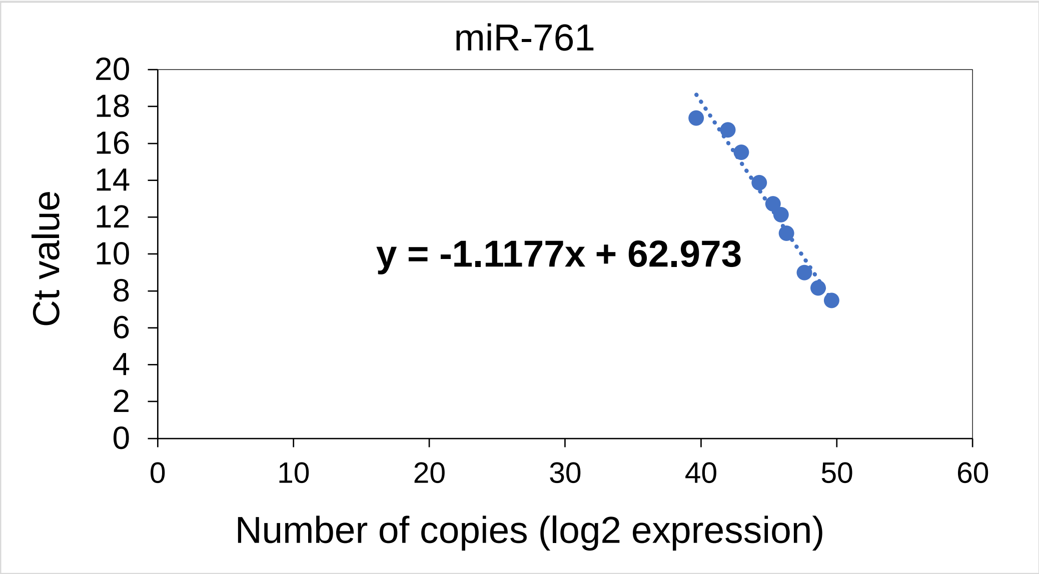


D

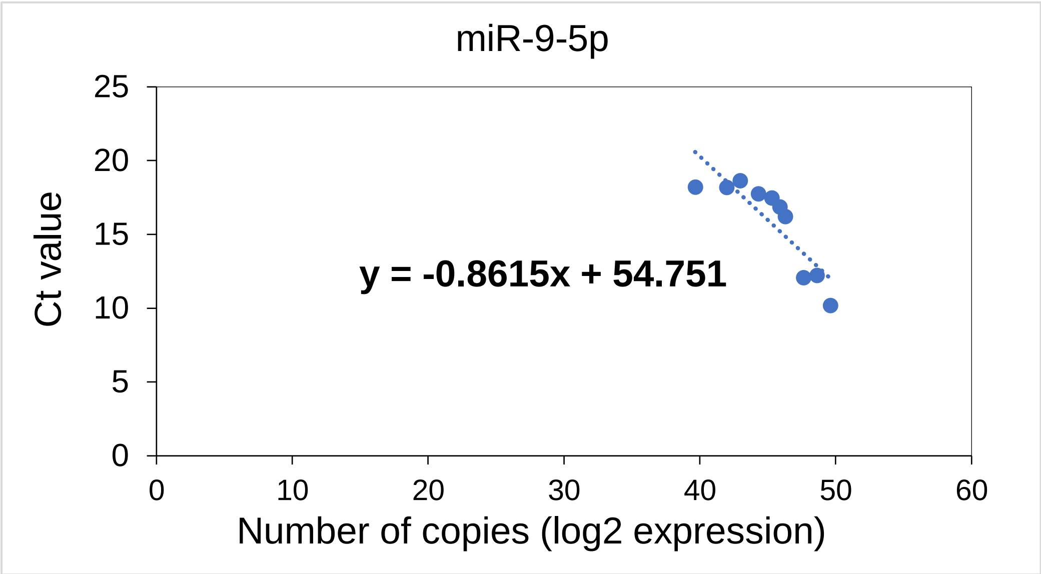


B

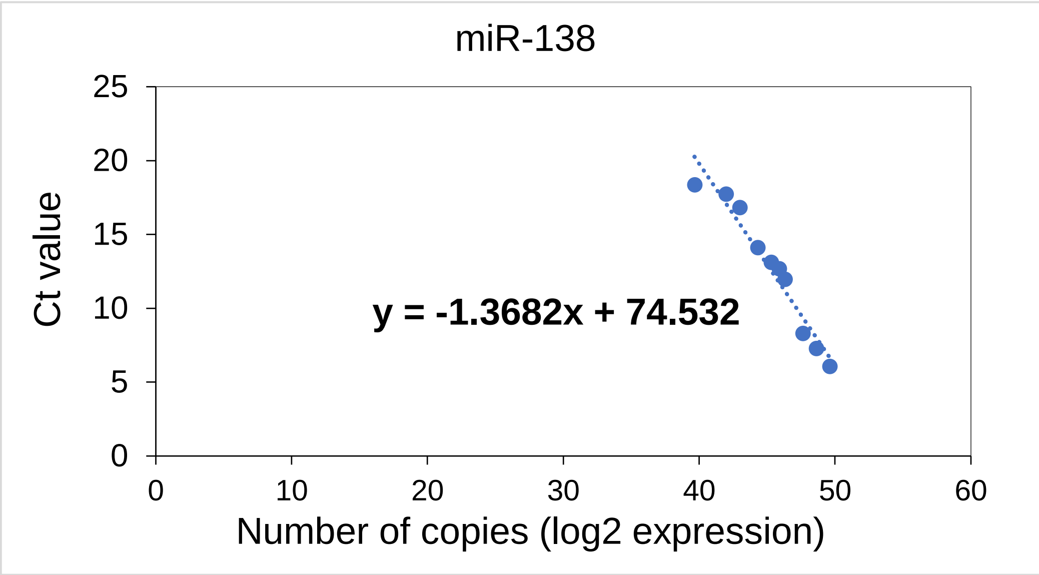


E

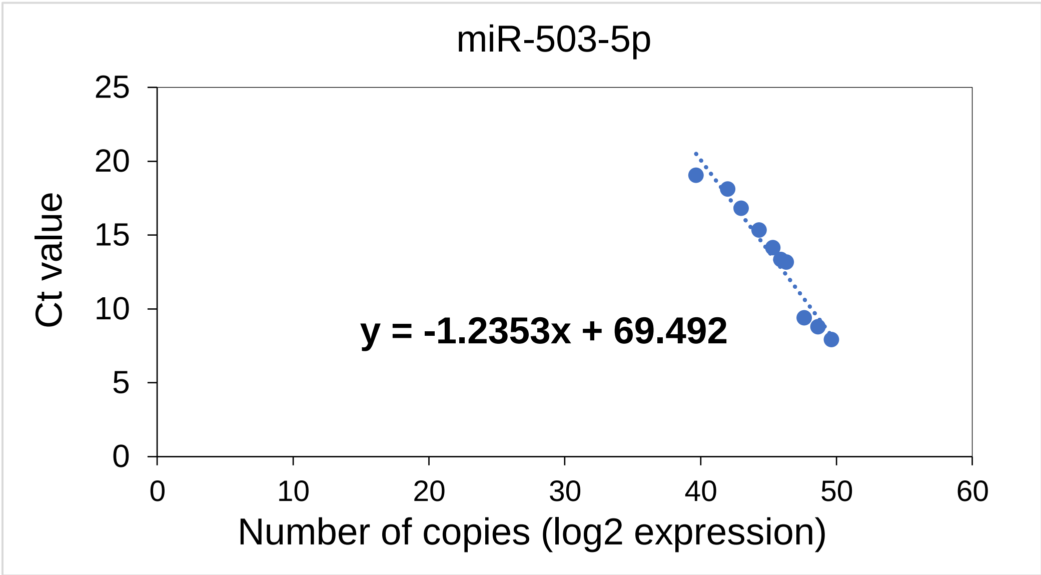


C

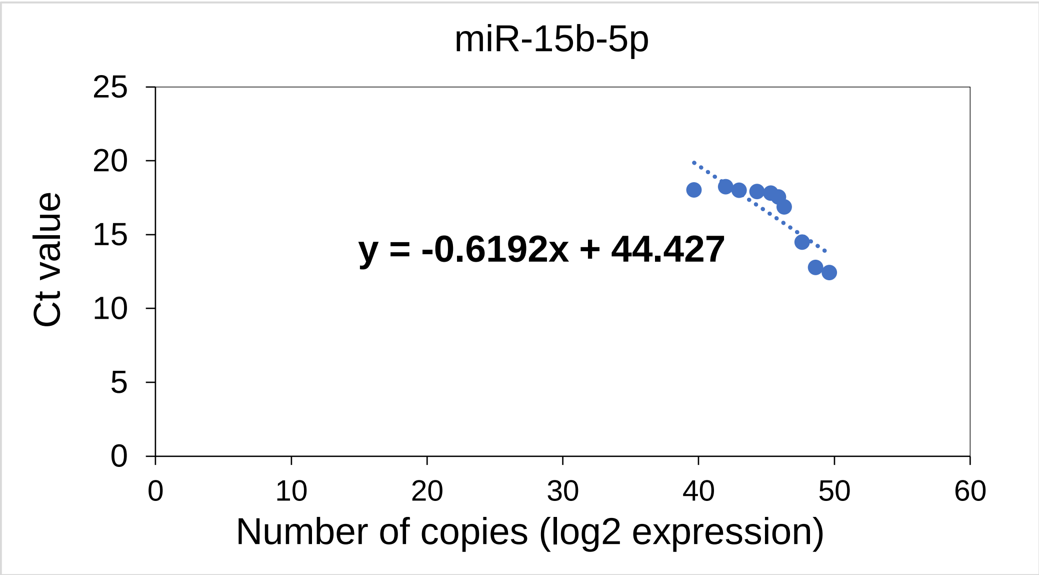


F

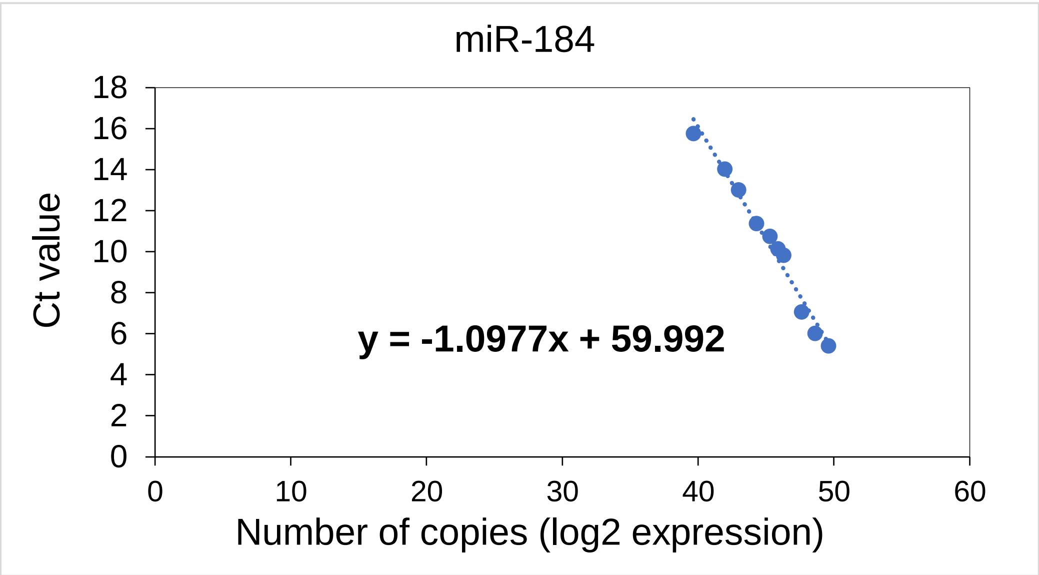


G

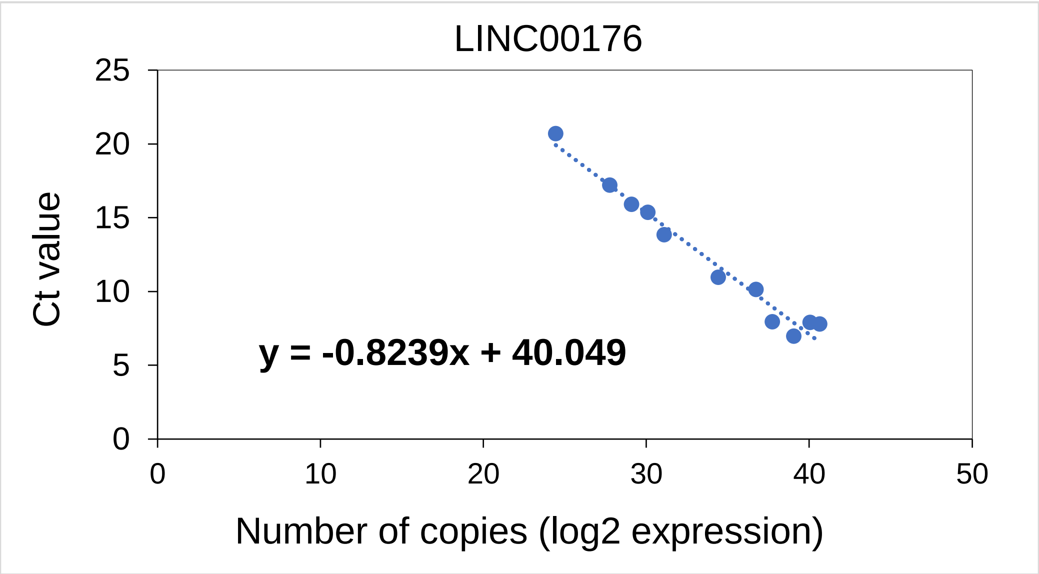


H

| RNA | Amount of RNA used for endogenous expression | Ct Value | Copy Number (log2 expression) |
|---|---|---|---|
| LINC00176 | 3 ug | 25.33 | 19.05 |
| miR-761 | 50 ng | 21.68 | 38.75 |
| miR-138 | | 20.74 | 46.16 |
| miR-15b-5p | | 20.24 | 31.74 |
| miR-9-5p | | 21.80 | 35.97 |
| miR-503-5p | | 23.95 | 39.91 |
| miR-184 | | 19.43 | 38.66 |
